# IL-16 production is a mechanism of resistance to BTK inhibitors and R-CHOP in lymphomas

**DOI:** 10.1101/2025.05.07.652612

**Authors:** Alberto J. Arribas, Francesca Guidetti, Eleonora Cannas, Luciano Cascione, Sara Napoli, Giulio Sartori, Federica Fuzio, Emma Pesenti, Chiara Tarantelli, Filippo Spriano, Antonella Zucchetto, Francesca M. Rossi, Alessio Bruscaggin, Andrea Rinaldi, Manuel Castro de Moura, Sandra Jovic, Andrea Raimondi, Roberta Bordone Pittau, Lodovico Terzi di Bergamo, Xiaofei Ye, Anastasios Stathis, Yaacov Ben-David, Qiang Pan-Hammarström, Federico Simonetta, Georg Stussi, Emanuele Zucca, Valter Gattei, Jennifer R. Brown, Manel Esteller, Davide Rossi, Francesco Bertoni

## Abstract

Introducing Bruton’s tyrosine kinase (BTK) inhibitors has significantly improved outcomes for patients with B-cell malignancies and autoimmune disorders. However, resistance, either primary or acquired, remains a major clinical challenge. To better understand the underlying resistance mechanisms to BTK inhibitors, we established an ibrutinib-resistant model from a patient-derived splenic marginal zone lymphoma (MZL) cell line (VL51) through prolonged drug exposure. Resistant cells exhibited a 15-fold increase in ibrutinib’s IC50, along with distinct morphological changes, mitochondrial activation, and cross-resistance to covalent, non-covalent BTK inhibitors and BTK degraders. Integrated transcriptomic, epigenomic, and proteomic analyses identified overexpression and secretion of IL-16 as a key feature of resistance, driven by chromatin remodeling and activation of the FLI1 transcription factor. IL-16 conferred ibrutinib resistance via CD9-mediated activation of the NF-κB and AKT signaling pathways and was found to be elevated in the serum of ibrutinib-refractory CLL patients. Functional studies showed that targeting the IL-16/CD9 axis using neutralizing antibodies or CD9-binding peptides restored sensitivity to BTK inhibitors and R-CHOP chemotherapy in MZL, mantle cell lymphoma, and diffuse large B-cell lymphoma models. These findings reveal a novel, targetable resistance mechanism with potential therapeutic implications for overcoming BTK inhibitor resistance in B-cell lymphomas.

## Introduction

Introducing pharmacological inhibitors of Bruton’s tyrosine kinase (BTK) in clinical practice has represented one of the most successful advances in treating patients with lymphoid tumors and autoimmune disorders ^1-4^. BTK is downstream of multiple receptors expressed and active in normal and neoplastic B cells, such as the B cell receptor (BCR), chemokine receptors (i.e., CXCR4), and Toll-like receptors. Covalent and non-covalent inhibitors, and, more recently, proteolysis targeting chimeras (PROTACs), have shown strong preclinical and clinical activity in patients with chronic lymphocytic leukemia (CLL), follicular lymphoma, mantle cell lymphoma (MCL), activated B-cell-like diffuse large B cell lymphoma (ABC DLBCL), and marginal zone lymphoma (MZL) ^1,2,5^.

Although BTK inhibitors have transformed the treatment landscape for B-cell malignancies, they are not curative. Some patients exhibit primary resistance, while others relapse or progress despite initial responses ^1,3,6^. We and others have reported genetic and non-genetic resistance mechanisms ^1,3,7-9^, but these findings do not fully account for the observed high inter- and intra-patient clinical variability. To further elucidate resistance mechanisms, we developed an ibrutinib-resistant cell line derived from a patient with splenic MZL, harboring somatic mutations in NOTCH2 and KLF2, hallmarks of the “NNK” (NF-κB/NOTCH/KLF2) splenic MZL cluster ^10^. This model identified interleukin-16 (IL-16) as a key driver of acquired resistance, conferring survival advantages through the CD9-dependent activation of NF-κB and PI3K-AKT signaling.

## Materials and Methods

### Cell lines culture

Cell lines were cultured according to the recommended conditions, as previously described ^11,12^. All media were supplemented with fetal bovine serum, penicillin-streptomycin-neomycin (5,000 units penicillin, 5 mg streptomycin, and 10 mg neomycin/mL, Sigma), and L-glutamine (1%). Cell lines were exposed to the IC90 concentration of ibrutinib (15 µM, Selleckchem, TX, USA) until they acquired specific drug resistance (resistant). In parallel, cells were cultured under similar conditions without a drug (parental). As previously reported ^11^, biological replicates were created by splitting the resistant cells after 1 month from the development of resistance, keeping them separate for 6 months before performing further experiments. VL51 parental and VL51 ibrutinib-resistant cells were stably transduced with mCherry as previously described ^13^ and used for the real-time cell growth experiments in the Incucyte S3 live-cell analysis instrument (Sartorius). Cells were periodically tested for their identity by short tandem repeat (STR) DNA profiling ^12^ and for Mycoplasma negativity ^11^.

### Transmission Electron Microscopy

Cell pellets were fixed with 2,5% glutaraldehyde in 0,1M cacodylate buffer, pH 7.4, for 1 hour at room temperature. After several washes in cacodylate buffer, samples were postfixed with reduced osmium solution (1% OsO4, 1,5% potassium ferrocyanide in 0,1M cacodylate buffer pH 7.4) for 2 hours on ice. After several washes in milli-Q water, sections were incubated in 0,5% uranyl acetate overnight at 4°C. Samples were then dehydrated with increasing concentration of ethanol, rinsed in propylene oxide, and infiltrated in a mixture of propylene oxide/epoxy resin (Epoxy Embedding Medium kit, Sigma-Aldrich) overnight at room temperature. Samples were finally embedded in pure resin and baked at 60°C for 48 hours. Ultrathin sections were obtained using an ultramicrotome (UC7, Leica microsystem, Vienna, Austria), collected on copper or nickel grids, stained with uranyl acetate and Sato’s lead solutions, and observed in a Transmission Electron Microscope Talos L120C (FEI, Thermo Fisher Scientific) operating at 120kV. Images were acquired using Velox software (FEI, Thermo Fisher Scientific). For morphological analysis, cellular protrusions and mitochondria were segmented using the segment-anything model tool in Microscope Image Browser software ^14^.

### Treatments

Ibrutinib, zanubrutinib, acalabrutinib, and vincristine were purchased from Selleckchem (Houston, TX, USA). Pirtobrutinib and NX-5948 were purchased from MedChemExpress (Monmouth Junction, NJ, USA). Human recombinant IL-16 was purchased from Prospec (Rehovot, Israel). Anti-IL-16 blocking antibody was purchased from BioLegend (anti-mouse IL-16 Functional Grade Purified neutralizing antibody, cat #519102). CD9 binding peptide (RSHRLRLH) as described by Suwatthanarak et al. ^15^ was synthesized by GeneSript (Rijswijk, Netherlands).

Response to single or drug combination treatments was assessed after 72 hours of exposure to increasing doses of drug followed by MTT assay. Sensitivity to single drug treatments was evaluated by IC50 (4-parameters calculation upon log-scaled doses, PharmacoGX R package ^16^) calculations. The beneficial effect of the combinations versus the single agents was considered both as synergism according to the Chou-Talalay combination index ^17^, the MuSyC algorithm ^18^, and to the highest single agent (HSA) and zero interaction potency (ZIP) synergy models included in the SynergyFinder R package ^19^.

### Flow cytometry and immunoblotting

Surface expression of CD4, CD9, and CXCR3 was measured by flow cytometry as previously described ^20^, using the antibodies listed in Table S1. Cells that underwent intracellular staining were spun down and washed twice in cold PBS to avoid signal from secreted IL16. Cells were then fixed, permeabilized, and stained with anti-IL16 antibody (Table S1). Finally, expression of intracellular IL16 was measured by flow cytometry. Protein expression levels were analyzed by flow cytometry with the flowCore R package ^21^. Immunoblotting was performed to determine the expression of AKT/p-AKT, NFκB1, NFκB2, RELA, RELB, and vinculin, as previously described ^22^, using the antibodies listed in Table S2.

### Enzyme-linked immunosorbent assay (ELISA)

ELISA (Human Cytokine Array Panel A, R&D Systems) was performed according to the manufacturer’s protocols, on conditioned media processed as previously described ^11^. ELISA on frozen human serum samples was performed using Luminex Assay (R&D Systems) according to the manufacturer’s protocols. The serum samples were collected from patients treated with ibrutinib and enrolled in tissue banking protocols at Dana-Farber Cancer Institute; all patients signed written informed consent before a sample was drawn, and the protocols were approved by the Dana-Farber Harvard Cancer Center Institutional Review Board.

### Genomics

For whole-exome sequencing (WES), libraries from genomic DNA were made with the exome capture SureSelect XT library preparation (v6, 58 Mb; Agilent Technologies) and sequenced using the HiSeq 2500 analyzer (Illumina) with the paired-end 2×125 bp read option to obtain a 14x coverage, as previously described ^11^. Transcriptome sequencing (RNA-Seq) was done using the TruSeq RNA Sample Prep Kit v2 for Illumina (Illumina, San Diego, CA, USA) as previously described ^11^. For small RNA-Seq, cDNA libraries were assembled using total RNA prepared using the SMARTer smRNA-Seq Kit for Illumina (Clontech Laboratories, Inc. USA), as previously described ^11^. Methylation profiling was done using the MethylationEPIC BeadChip Infinium following the manufacturer’s instructions for the automated processing of arrays with a liquid handler (Illumina Infinium HD Methylation Assay Experienced User Card, Automated Protocol 15019521 v01), as previously described ^11^.

Targeted sequencing was done using the LyV4.0 CAPP-seq gDNA Assay with libraries generated with the KAPA Hyper Prep Kit (KAPA Biosystems) enriched using SeqCap HyperChoice Library probes (NimbleGen; Roche Diagnostics, Jakarta, Indonesia). Libraries were sequenced on the NextSeq500 (Illumina, San Diego, CA) instrument by paired-end sequencing (2 × 150-cycle protocol), as previously described ^23^.

### Assay for Transposase-Accessible Chromatin with sequencing (ATAC-seq)

ATAC-seq for cell lines was performed following the Omni-ATAC-seq protocol ^24^ with some modifications. Fifty thousand viable cells were lysed in 10 mM tris-HCl (pH 7.4), 10 mM NaCl, 3 mM MgCl2, and 0.1% Igepal CA-630 (Nonidet P-40). The nuclear pellet was then subjected to transposition reaction in 50 μL volume using Illumina Tagment DNA Enzyme and Buffer Small Kit (catalogue n. 20034197) at 37°C for 30 minutes. Then 9 μL per well of stopping mix was added (Clean up buffer 5 μL (900 mM NaCl and 30 mM EDTA), 5% SDS 2 μL and Proteinase K (20 mg/ml) 2 μL) and then incubated at 40°C for 30 min. DNA was then cleaned up with Agencourt AMPure XP-Beckman Coulter. PCR amplification was performed according to the protocol for OneTaq Hot Start 2X Master Mix with GC Buffer (M0485, New England Biolabs). The libraries were cleaned up with Agencourt AMPure XP-Beckman Coulter.

### Data mining

Data mining of WES, RNA-Seq, small RNA-Seq, methylation profiling and cytokine array data was performed as previously described ^11^. The false discovery rate (FDR, Benjamin-Hochberg correction) was calculated to control for false positives. FDR <0.05 and absolute fold-change higher than 2 (RNA-seq) or 0.3 (corresponding to 30% of differences in Beta fold-change or in normalized fold-change from the cytokine array) was considered significant. Functional analysis was performed on the collapsed gene symbol list using GSEA (Gene Set Enrichment Analysis) with the MSigDB (Molecular Signatures Database) C2-C7 gene sets ^25^, and SignatureDB database (https://lymphochip.nih.gov/signaturedb/). Statistical tests were performed using the R environment (R Studio console; RStudio, Boston, MA, USA). Finally, unsupervised multidimensional scaling plots were used to visualize resistant and parental multi-omics profiles.

Data mining of targeted DNA sequencing was performed as previously described ^23^.

For ATAC-seq, the Nextflow-based pipeline “atacseq” was used (https://nf-co.re/atacseq, v.2.1.1). Briefly, reads are trimmed using the TrimGalore! tool and aligned to the GRCh38 assembly of the human genome using BWA (v. 0.7.17-r1188). Duplicate reads are marked using Picard (v.3.0.0). Reads mapping to mitochondrial DNA and to ENCODE blacklisted regions were removed. Genome browser tracks are created with the genomeCoverageBed command in BEDTools (v. 2.30.0) and normalized such that each value represents the read count per base pair per million mapped. Finally, the UCSC Genome Browser’s bedGraphToBigWig tool (v. 445) is used to produce a bigWig file. Peak calling was performed using MACS2 (v.2.2.7.1.0) using the ‘-nomodel’ parameter and reads counted for each sample in each consensus peak using featureCounts of the subread package (v. 2.0.1) to create a matrix of count for each sample and each peak region. The list of consensus peaks was obtained by merging all peaks in all samples by bedtools merge (v. 2.30.0). Consensus peaks were mapped to genomic location, including gene and distance from the nearest Transcription Start Site (TSS) using the ChIPpeakAnno and the BiomaRt packages in R. Read counts were normalized across samples using the quantiles normalization function ‘normalize.quantiles’ from the preprocessor package in R. Inspection of batch effect artifacts were exploited with Principal Component Analysis (PCA). Supervised differential accessibility was performed using DeSeq2 package in R. Region-set enrichment analysis was performed with the Locus Overlap Analysis (LOLA) tool ^26^.

## Results

### Development of ibrutinib-resistant cells in marginal zone lymphoma

VL51 cells were exposed to increased concentrations of ibrutinib for six months until the acquisition of stable resistance. The developed ibrutinib-resistant model (RES) had a 15-fold higher IC50 than parental cells (PAR).

Calculation of IC50 was validated after three weeks of culture in a drug-free medium. Resistant cells exhibited decreased sensitivity to covalent and non-covalent BTK inhibitors and BTK degraders (Figure S1). Multidrug resistance phenotype was evaluated by maintaining sensitivity to the MDR-substrate vincristine and no increased expression levels of MDR1/2 (Figure S2).

We then studied potential changes in the morphology of VL51 ibrutinib-resistant cells using electron microscopy compared to their parental counterpart. Ibrutinib-resistant cells were smaller than parental cells but with more filopodia/membrane protrusions (Figure S3A). In addition to the more complex membrane, the mitochondria were rounder, bigger, and more fragmented in resistant than in parental cells, suggesting a higher mitochondria activation in RES (Figure S3B).

Finally, resistant cells also exhibited no significant changes in sensitivity to compounds targeting downstream BCR signaling, including PI3K inhibitors (idelalisib, duvelisib, copanlisib, and umbralisib) (Figure S4).

### Upregulation and secretion of IL-16 in VL51 IBR-RES cells

No DNA copy number variations or mutations were detected by WES or by targeted sequencing for genes recurrently mutated in lymphomas (including *BTK* and *PLCG2)* in resistant cell lines compared to parental cells (Table S3), indicating a non-genetic mechanism of resistance. Despite the similar DNA profiling, resistant cells showed distinctive transcriptomic and methylation profiles from the parental counterparts (Figure 1A). At the transcriptome level, RES exhibited overexpression of genes coding for proteins involved in cytokine secretion (*IL16, LCP2, CXCL10*), immune response (*FCER1A, CD300, CSF2*), cell proliferation (*CCNB1, MKI67*), integrins (*ITGAM, ITGA11*), and members of the NF-κB (*TNF, LTA, CD44*), PI3K-AKT (*PIK3R6, GPER1, PRR5L*) and genes related to resistance to targeted therapies in lymphoma (*EGR1/2, IGF1, BCL2A1*). Also, RES presented lower expression of tumor suppressor genes (*CDKN2B, CDKN2A, CDKN1A*), negative regulators of PI3K-AKT-mTOR (*PIK3IP1, DEPTOR*), genes located at 7q31, frequently deleted in SMZL (*TASR3, CAV1, DNAJB9*) ^27,28^ and apoptosis-related genes (*PDCD4, ANXA1*) (Figure 1A, Table S4).

**Figure 1.**
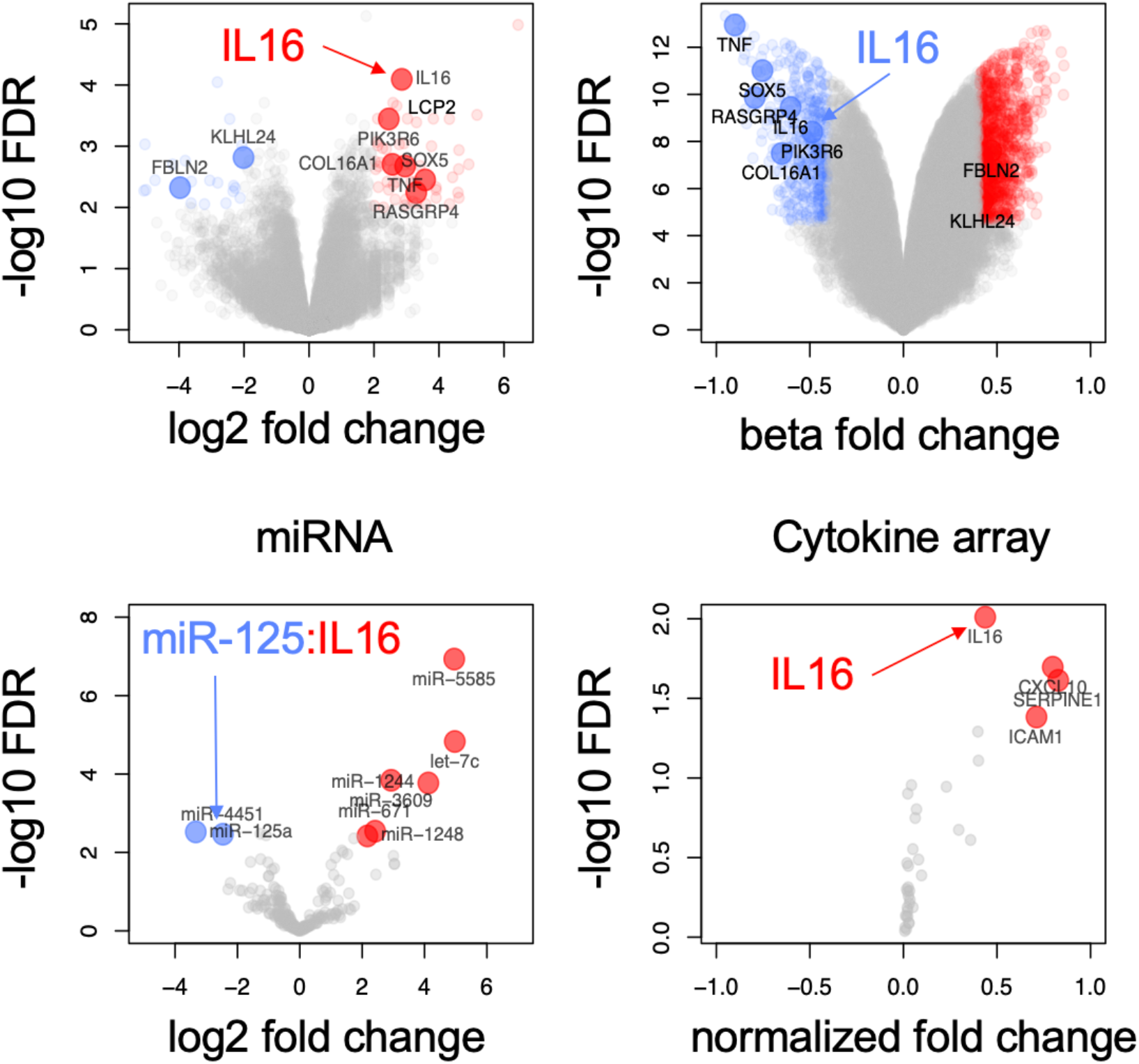
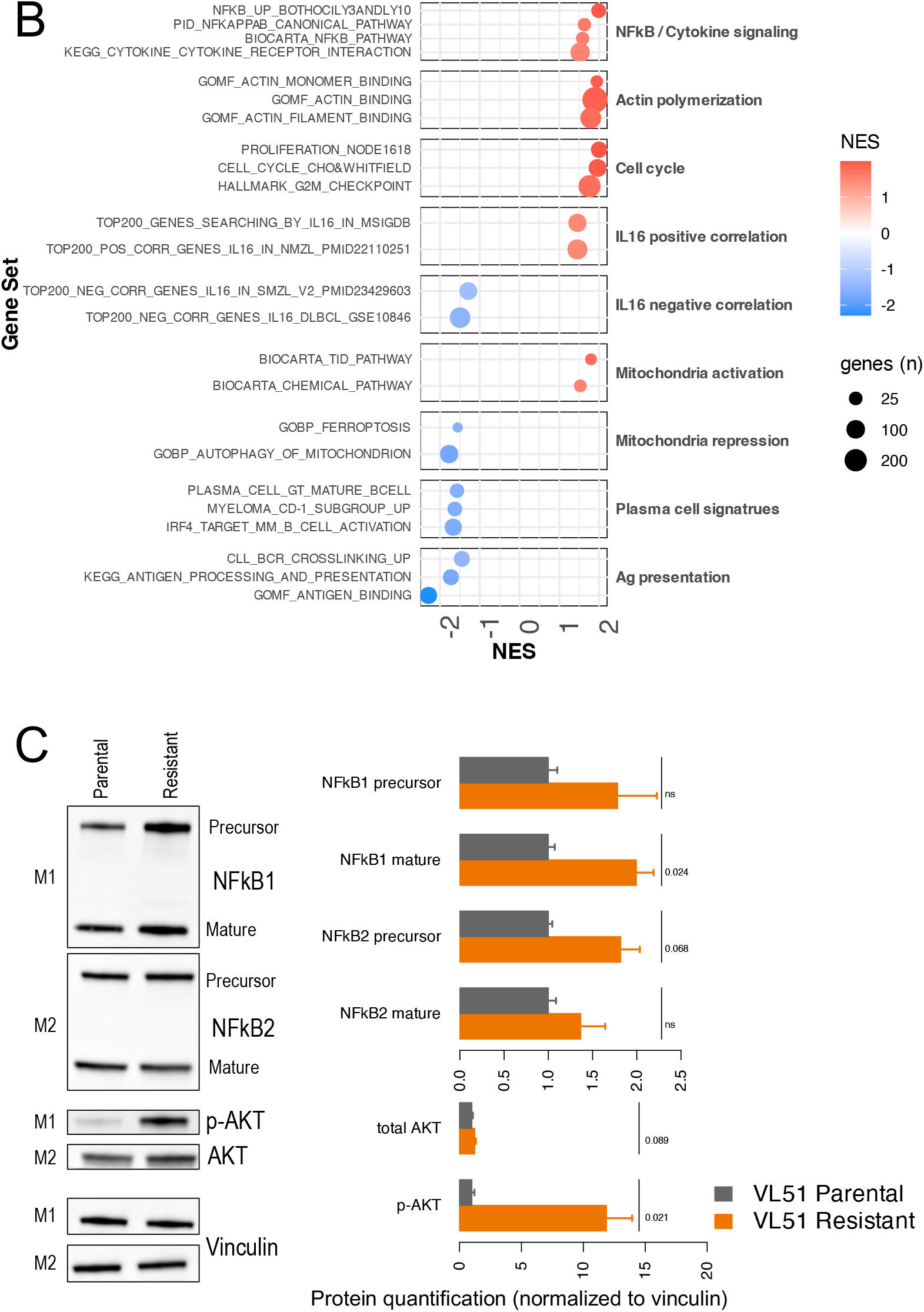
Multi-omics signature of resistance identifies overexpression and secretion of IL16. (A) Vulcano plot of RNA (GEP), methylation, miRNA, and secretome (cytokine array) profiles are resistant compared to parental. Vulcano plots represent the fold change between resistant and parental in the x-axis (log_2_ fold change for RNA and miRNA, Beta-value fold change for methylation, and normalized fold change for cytokine array) and -log_10_ FDR (false discovery range, Benjamin-Hochberg correction) in the y-axis. Values from a moderated t-test. Genes up-regulated and hypomethylated are highlighted in red for resistant and in blue for parental. (B) Top enriched genesets in resistant (red) and parental (blue) by Gene set enrichment analyses (GSEA). (C) Protein levels of NFkB1 or NFkB2 (precursor and mature forms) and total or phospho-AKT by immunoblot in parental (grey) and resistant (orange) cells. Values correspond to the average of protein quantification normalized to vinculin levels in two independent experiments (the second experiment in Fig S8). M1 and M2 correspond to the two immunoblot membranes used for the different antibodies. Error bars represent the standard deviation of the mean. P values from non-parametric t-test, statistically significant for p< 0.05.

Integration of transcriptome and DNA promoter methylation profiles of RES and PAR cells identified up-regulated and unmethylated genes, including secreted factors such as IL16 and TNF, and PI3K-AKT signaling genes (PIK3R6, RASGRP4). Comprehensive analyses on small RNA and RNA profiles identify up-regulated miRNAs (miR-5585, let-7c, miR-1244) and two repressed miRNAs that target IL-16: miR-125a and miR-4451 ^29^ (Fig 1A, Table S4). Gene-set enrichment analyses showed enrichment of genes involved in NF-κB signaling, actin polymerization, cell cycle, and IL-16-associated signatures in RES. In contrast, PAR cells were characterized by elevated antigen presentation and IRF4-induced/plasma cell signatures (Figure 1B). Consistent with increased activity of the mitochondria in RES, the latter presented an enrichment of genes belonging to pathways related to mitochondria activation, and a downmodulation of genesets related to the repression of the mitochondria (Figure 1B).

Based on the upregulation of secreted factors observed at RNA level, we tested the sensitivity to ibrutinib of the VL51 parental cells cultured with the resistant-conditioned medium. Consistent with the role of secreted factors in the resistance mechanism, we observed decreased sensitivity to ibrutinib in parental cells cultured for 48 hours in the resistant-conditioned medium (Figure S5). Next, we investigated the secretome profile on the conditioned media (72 hr) of PAR and RES by cytokine array, which confirmed the secretion of IL-16 and CXCL10 (Figure 1A, Figure S6, Table S4). In accordance with the *in vitro*, secretome and RNA-Seq findings, RES exhibited higher intracellular IL-16 expression than PAR (Figure S7).

At immunoblotting, we observed activation of NF-κB and strong pAKT upregulation in resistant VL51 cells (Figure 1C, Figure S8).

### IL-16 induces resistance to ibrutinib in lymphoma cells

Next, we investigated whether stimulation with IL-16 may reduce the response to ibrutinib. Signaling driven by IL-16 relies on the binding to two potential surface receptors: CD4 or CD9 ^30^. Since the T-cell marker CD4 is not expressed in B-cells, as also confirmed at the RNA level in our models, we investigated the surface expression of CD9 in the VL51 parental and resistant cells, where CD9 was well expressed in both, with slightly higher expression in the resistant cells (Figure S9). Recombinant IL-16 induced resistance to ibrutinib in VL51 parental cells with limited effect in resistant cells, consistent with the constitutive secretion of IL-16 in the latter (Figure 2A-B). Then, to further investigate the impact of IL-16 in the response to ibrutinib, we stimulated two CD9-positive B-cell lymphoma models, selected taking advantage of previous transcriptome data we had obtained ^31^, with recombinant IL-16. We observed a reduction in the sensitivity to ibrutinib in MEC1 (CLL) and REC1 (MCL) cells (Figure 2C, Figure S10).

**Figure 2.**
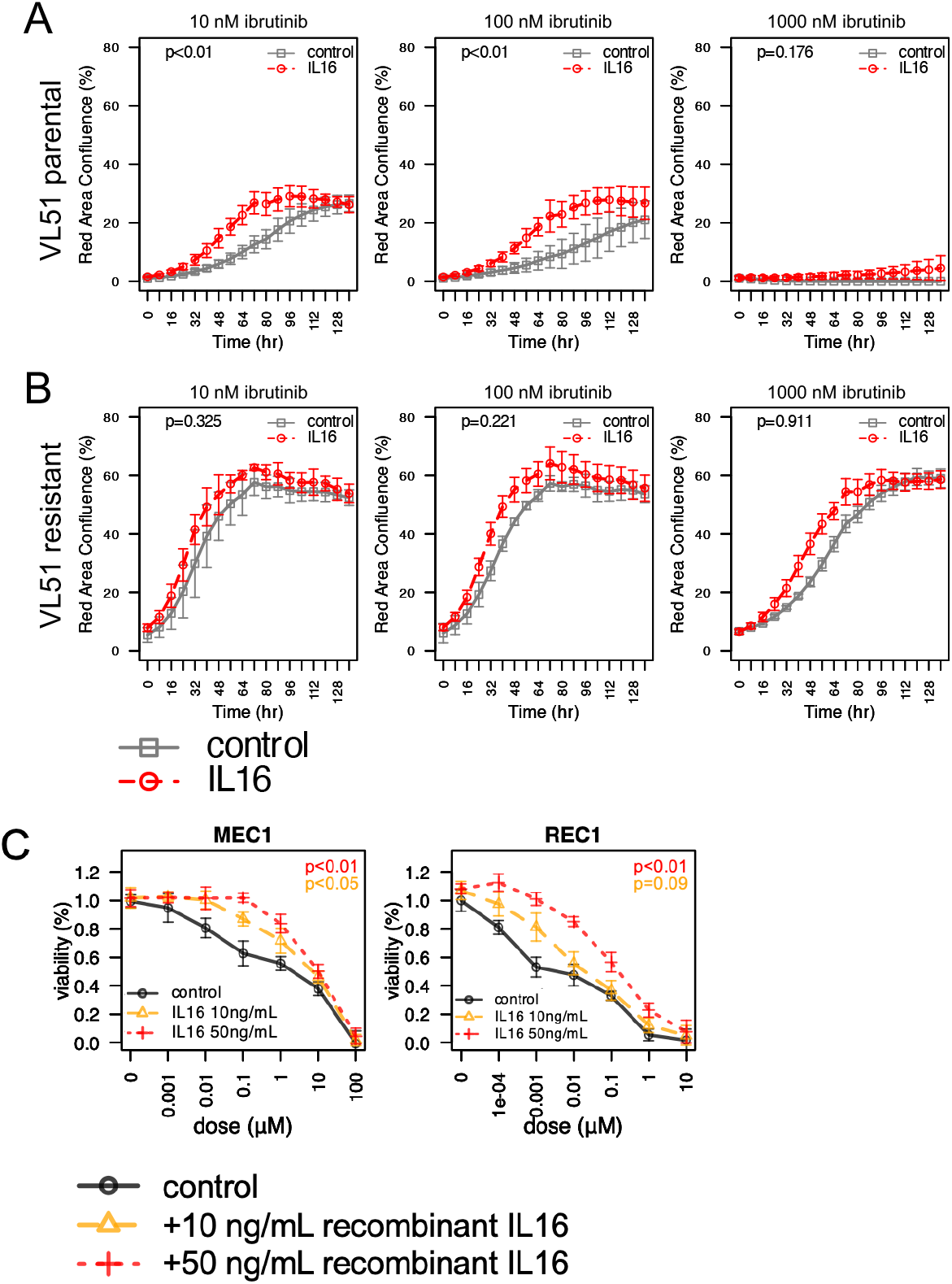
Human recombinant IL-16 induces resistance to ibrutinib. VL51 parental (A) or ibrutinib-resistant (B) cells with stable expression of mCherry were cultured for 6 days in the presence of recombinant IL-16 (30ng/mL, red) or PBS (control, grey) under exposure to increasing concentrations of ibrutinib: 10 nM (left), 100 nM (center) or 1 µM (right). Two independent experiments were performed using the Incucyte S3 live-cell analysis instrument. Cell growth was estimated by the signal of mCherry expression (red channel) measured by the percentage of red area confluence. P represents the p-value from the Z-test comparing IL-16 vs control (PBS). (C) MEC1 (left) or REC1 (right) cells were cultured for 72 hours in the presence of recombinant IL-16 (10ng/mL in yellow, or 50ng/mL in red) or PBS (control, black) under exposure to increasing concentrations of ibrutinib, followed by MTT assay. Three independent experiments were performed. P represents the p-value from the Z-test comparing each IL-16 dose vs control (PBS).

### IL-16 is enriched in the serum of CLL patients who do not respond to ibrutinib

To investigate the clinical relevance of our findings, secreted levels of IL-16 were evaluated in the serum of CLL patients with resistance to the BTK inhibitor in the absence of BTK or PLCG2 mutations, compared to patients responding to the drug and paired for clinical features. In agreement with our hypothesis, serum from resistant patients exhibited significantly higher levels of IL-16 compared to responders (Figure 3).

**Figure 3.**
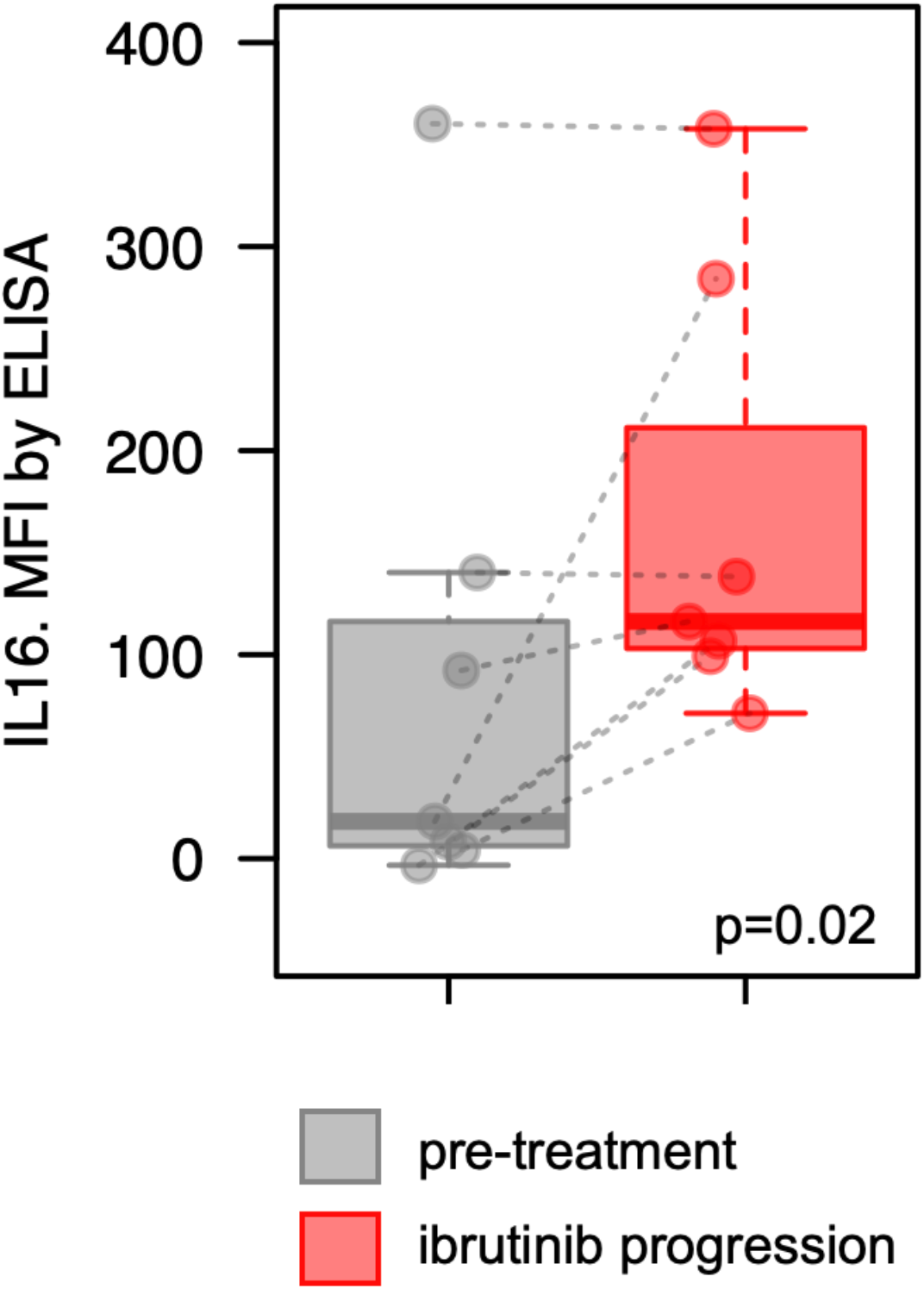
Higher IL-16 levels are detected in serum from ibrutinib-resistant CLL patients. IL-16 secretion was evaluated in the serum of CLL samples at the end of the treatment by ELISA (Luminex, R&D Systems). (A) IL-16 secretion in the serum of patients treated with ibrutinib was compared at the time of progression (red) with baseline levels of ibrutinib treatment (pre-treatment, grey). P for p-value from a non-parametric t-test. MFI: mean fluorescence intensity.

### IL-16-driven gene expression profiling and signaling activation

Hence, we investigated the transcriptome changes orchestrated by IL-16 in VL51 parental cells. VL51 parental cells were stimulated with 30 ng/ml for 24 hours and underwent RNA-Seq. Upon exposure to IL-16, VL51 parental cells exhibited enrichment in BCR-NF-κB and NOTCH signaling pathways, MYC targets, and unfolded protein response while downmodulating oxidative phosphorylation (OxPhos) and Interferon cascade (IFN). The top upregulated genes included NF-κB-induced chemokines (*CCL3/4*), phosphatases (*DUSP2/4*), ETS transcription factors (*ETV4/5*), and genes involved in ibrutinib resistance (*EGR1/2/3*). Conversely, the top downregulated genes included cell adhesion molecules (*PECAM1, VCAM1*), interleukin modulators (*FYB*), interactors of cytoskeleton activity (*MYO18B*), and negative regulators of mTOR (*DEPTOR*) (Figure 4).

**Figure 4.**
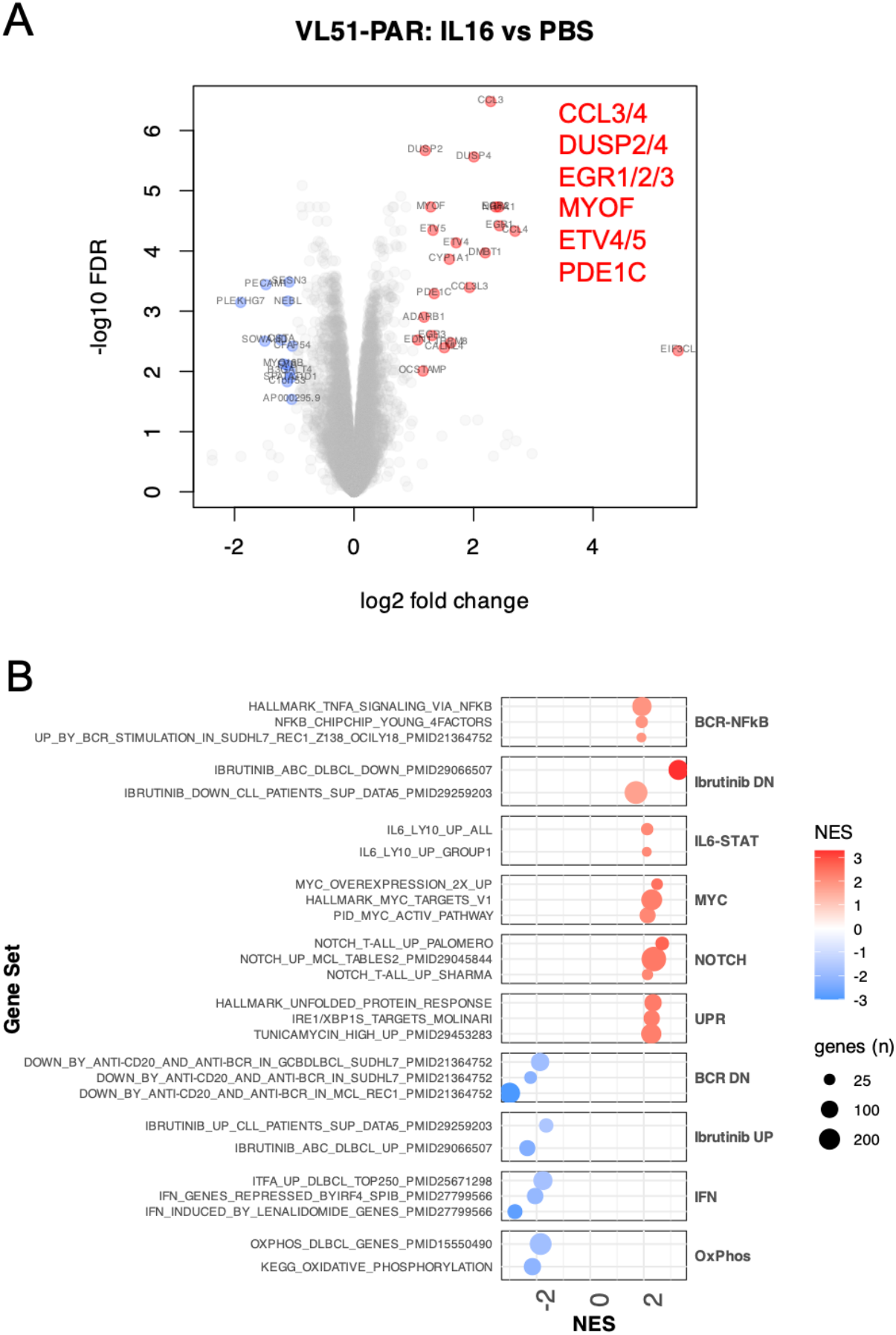
IL-16-driven transcriptome resembles the ibrutinib-resistant signature. The gene expression profile was evaluated by RNA-sequencing in VL51 parental cells, either stimulated with human recombinant IL-16 (30ng/mL) or PBS (control) for 24hr. (A) Vulcano plots represent the fold change between IL-16 and control in the x-axis (log_2_ fold change) and -log_10_ FDR (false discovery range, Benjamin-Hochberg correction) in the y-axis. Values from a moderated t-test. Red was used for overexpression, and blue was used for repression in IL-16-stimulated compared to control (PBS). (B) Top enriched genesets in IL-16 (red) and parental (blue) by Gene set enrichment analyses (GSEA).

Consistent with the role of IL-16 in resistance to ibrutinib, gene expression profile of RES showed enrichment of the signature of recombinant IL-16 (Figure S11), and recombinant IL-16 partially induced the ibrutinib-resistant signature, including genes related to resistance to targeted therapies (*IGF1, EGR1/2, BCL2A1*), NF-κB genes (*TNFRSF13B, NR4A1*), MYC targets (*ODC1, NOLC1, NOP16*), apoptosis (*PDCD4, ANXA1*), TNF signaling (*TNFAIP2, RNF19B*) and the negative regulators of PI3K-AKT-MTOR *DEPTOR* (Figure S12). IL-16 stimulation repressed genes involved in the negative regulation of actin polymerization, accordingly with the observed enrichment of actin polymerization signatures in VL51 ibrutinib-resistant cells (Figure S13). Notably, genes involved in mitochondrial biogenesis were upregulated in parental cells upon recombinant IL-16, while apoptosis was reduced by exposure to the cytokine (Figure S14).

Based on the gene expression profile of ibrutinib-resistant and the transcriptome orchestrated by IL-16 in parental cells, we investigated NF-κB activation in the VL5 lines. In agreement with the transcriptome data, ibrutinib-resistant cells exhibited up-regulation of NF-κB cascade, which recombinant IL-16 increased in VL51 parental cells (Figure S15).

Since CXCL10 is an NF-κB target, we wondered whether IL-16 may induce CXCL10 secretion upon NF-κB activation in ibrutinib-resistant cells. Using real-time PCR, we measured CXCL10 expression upon recombinant IL-16 in VL51 cells, where we observed an upregulation of CXCL10 by exposure to IL-16. Supporting a potential role of CXCL10 in ibrutinib resistance, its receptor CXCR3 was well expressed in VL51 cells (surface expression by FACS). Thus, we tested the effect of recombinant CXCL10 in the response to ibrutinib in VL51 parental cells, observing a decrease in the ibrutinib response upon CXCL10 stimulation (Figure S16).

Finally, we investigated whether these results might be extrapolated to different *in vitro* models. IL-16 and CD9 but not CXCL10 expression levels were inversely correlated with sensitivity to ibrutinib in a panel of 13 B-cell lymphoma cell lines, which we had previously treated with the BTK inhibitor (Figure S17) ^32^.

### IL-16/CD9 axis is expressed in B-cell lymphoma clinical specimens and confers R-CHOP resistance in ABC-DLBCL cells

First, taking advantage of public expression datasets of B-cell lymphomas ^33-36^, we investigated the potential relevance of the IL-16/CD9 axis in clinical specimens. We confirmed that both IL-16 and CD9 were expressed across non-Hodgkin lymphoma clinical specimens and in malignant cells of DLBCL specimens ^33-36^ (Figure S18). Moreover, since IL-16 is part of the activated B-cell-like (ABC) DLBCL transcriptome signature ^37-39^, we tested the effect of IL-16 stimulation on R-CHOP sensitivity in ABC DLBCL cell lines. OCILy10 and TMD8 cell lines, derived from ABC DLBCL, with high sensitivity to R-CHOP *in vitro* (IC50∼50nM), were tested for changes in drug response upon stimulation with recombinant IL-16. We observed that IL-16 stimulation decreased sensitivity to R-CHOP in ABC DBCL cell lines (Figure 5), while cell proliferation was not affected by the cytokine.

**Figure 5.**
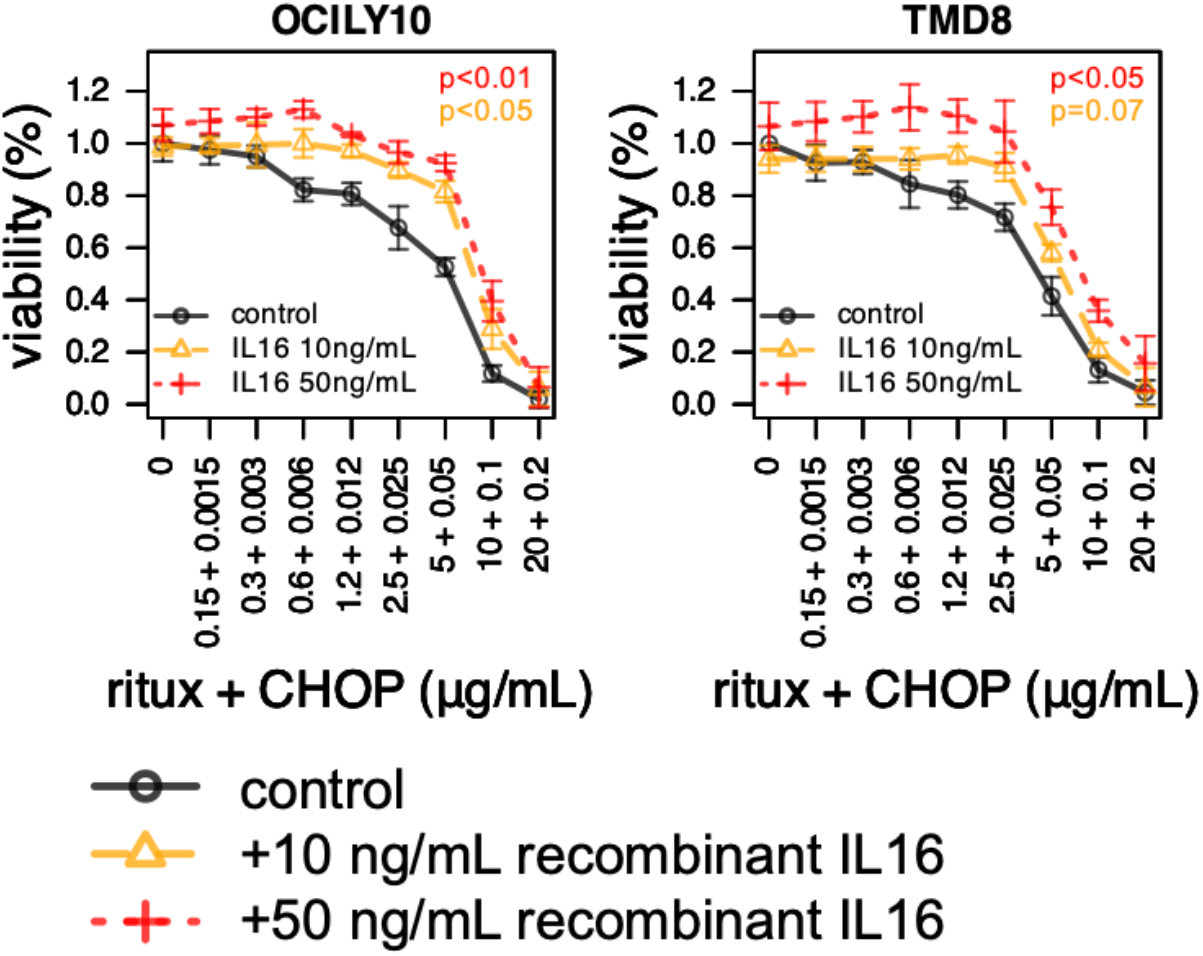
Human recombinant IL-16 induces resistance to RCHOP in ABC-DLBCL models. OCI-Ly10 (left) or TMD8 (right) cells were cultured for 72 hours in the presence of recombinant IL-16 (10ng/mL in yellow, or 50ng/mL in red) or PBS (control, black) under exposure to increasing concentrations of the *in vitro* equivalent of RCHOP, followed by MTT assay. Three independent experiments were performed. P represents the p-value from the Z-test comparing each IL-16 dose vs control (PBS).

### Pharmacological inhibition of the IL-16/CD9 axis improves the sensitivity of MZL and MCL cells to BTK inhibitors

We then explored the possibility of blocking the IL-16/CD9 axis using an anti-IL-16 neutralizing antibody ^40^ or an eight-mer peptide that binds and inhibits CD9 (CD9-BP) ^40^.

Thus, we exposed the RES cells and other CD9-positive cell lines with high expression of IL-16 and different degrees of sensitivity for the BTK inhibitors to ibrutinib or zanubrutinib as single agents plus the IL-16/CD9 targeting compounds (Figure S10). In the IL-16-positive SSK41 (MZL model) and MAVER1 (MCL model), and in accordance with the effect observed in VL51 PAR cells, upon IL-16 stimulation, the response to BTK inhibitors was attenuated, and conversely, the anti-IL-16 antibody or CD9-BP enhanced sensitivity to ibrutinib or zanubrutinib (Figure 6A-B). Additionally, the combination of ibrutinib and anti-IL-16 antibody was beneficial in MZL and MCL models *in vitro* (Figure 6A). As negative controls, we tested the CD9-low Karpas1718 and its ibrutinib-resistant derivative K1718-RES ^9^. No effect was seen upon IL-16 stimulation or CD9 blocking (Figure 6B).

**Figure 6.**
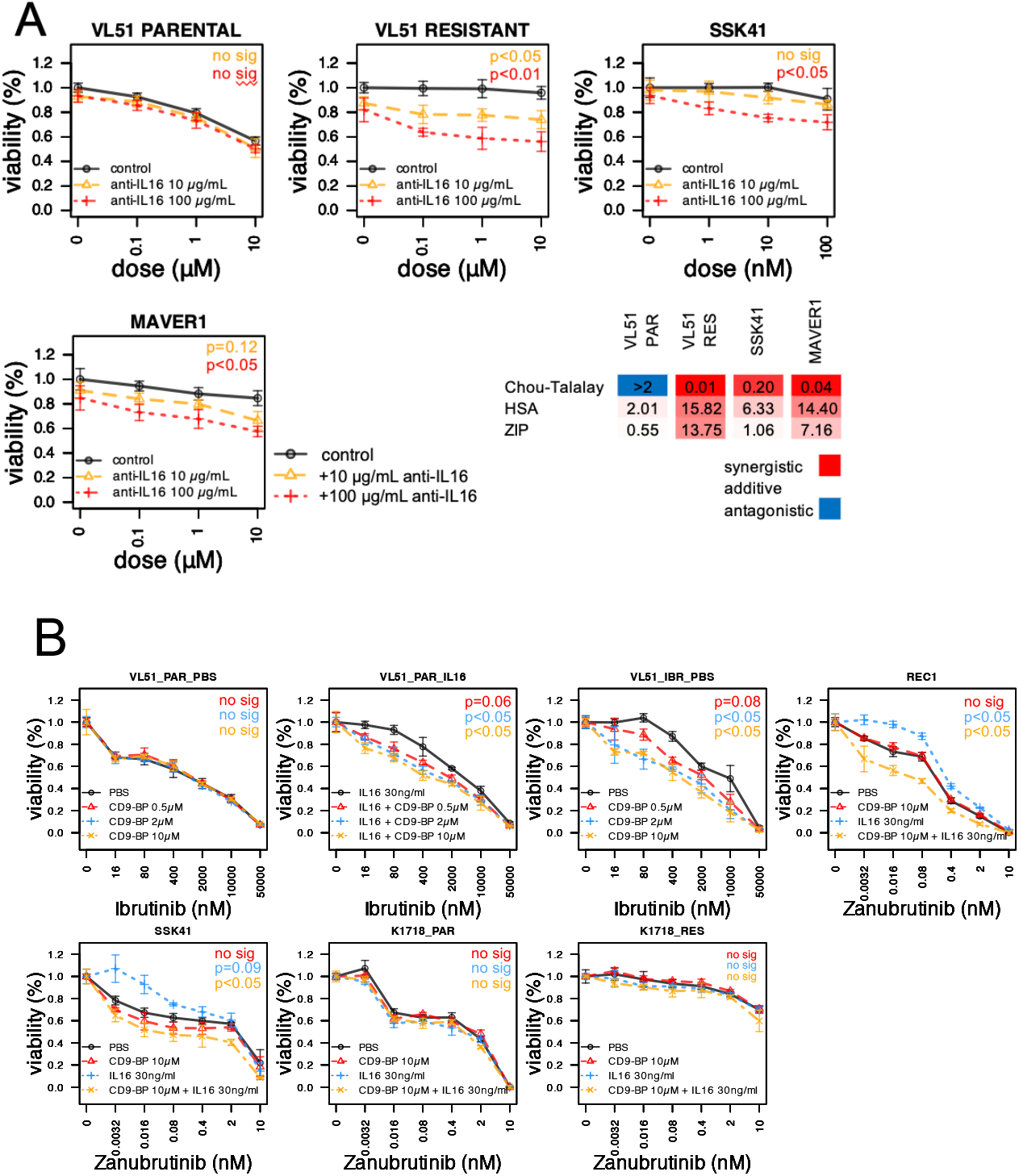
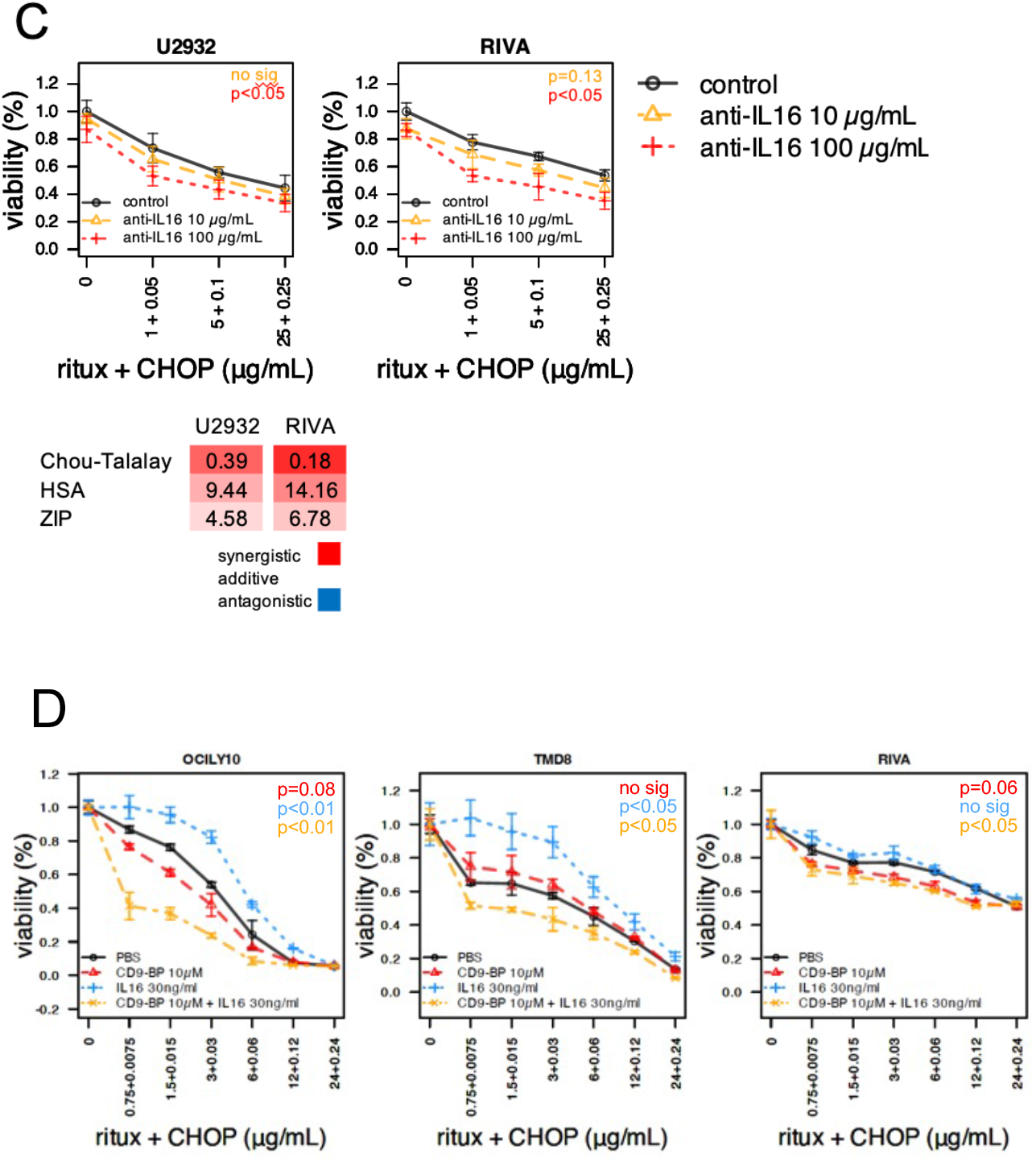
Blocking the IL-16-CD9 axis improves response to ibrutinib and RCHOP in B-cell lymphoma. (A) VL51 parental (IL-16-negative) and VL51 ibrutinib-resistant, SSK41 and MAVER1 (all IL-16-positive) cells were cultured for 72 hours in the presence of the blocking antibody for IL-16 (10µg/mL in yellow, or 100µg/mL in red) or PBS (control, black) under the exposure to increasing concentrations of ibrutinib followed by MTT assay. (C) U2932 (IL-16-negative) and VL51 ibrutinib-resistant, SSK41 and MAVER1 (all IL-16-positive) cells were cultured for 72 hours in the presence of the blocking antibody for IL-16 (10µg/mL in yellow, or 100µg/mL in red) or PBS (control, black) under the exposure to increasing concentrations of ibrutinib (A-B) or the *in vitro* equivalent of RCHOP (C-D), followed by MTT assay. Three independent experiments were performed. P represents the p-value from the Z-test comparing each IL-16 dose vs control (PBS).

Mechanistically and in accordance with the signaling observed in VL51-resistant cells, recombinant IL-16 induced NF-κB activation and increased pAKT in VL51 parental cells, which was abrogated by the addition of the CD9-binding peptide (Figure S19).

### Pharmacological inhibition of the IL-16/CD9 axis improves sensitivity to R-CHOP in ABC DLBCL cells

Based on the fact that IL16 is part of the ABC DLBCL signature, we then assessed the interference of IL-16/CD9 axis in the response to R-CHOP in four ABC DLBCL cell lines expressing IL-16/CD9 (U2932, OCI-Ly10, TMD8 and RIVA) (Figure S10). The blocking antibody anti-IL-16 improved the sensitivity to R-CHOP in U2932 and RIVA models, in which the combination was synergistic (Figure 6C). Similarly, when stimulated with IL-16, the addition of CD9-BP enhanced the response to RCHOP in OCI-Ly10, TMD8, and RIVA cell lines (Figure 6D). All together, these data suggest a role for IL16 in resistance not only to BTK inhibitors also but to first-line chemoimmunotherapy in ABC DLBCL. These findings support the therapeutic potential of targeting the IL-16/CD9 axis to overcome drug resistance and improve treatment efficacy in this aggressive lymphoma subtype.

### Chromatin remodeling facilitates IL-16 transcription through FLI1 transcription factor

Finally, we analyzed the chromatin accessibility of the VL51-resistant cells to understand further the epigenetic reprogramming underlying the resistance mechanism. The analysis revealed significant changes compared to the parental cell line. Specifically, we observed increased chromatin accessibility in regions associated with genes upregulated at the RNA level (Figure S20A). In contrast, regions with decreased chromatin accessibility corresponded to downregulated genes in the RNA-seq data (Figure S20B).

Consistently, both the promoter and intergenic regions of the IL-16 gene were among the most accessible regions in resistant cells, and the latter exhibited enrichment in the peaks at the IL-16 promoter compared to the parental cells (Figure 7A-B). We then performed region-set enrichment analysis to investigate the functional implications of epigenetic reprogramming. This analysis revealed increased activity of transcription factors involved in B cell development and function, including EBF1, PAX5, PU.1, and ETS family members such as FLI1 (Figure 7C). Notably, FLI1 induces the expression of genes involved in immune response and interleukin and BCR signaling ^41^. Indeed, FLI1 and its transcriptional signature were upregulated at the RNA level in the ibrutinib-resistant models (Figure S20C-D), and FLI1 expression correlates with Ibrutinib resistance in ABC-DLBCL models and with IL16 mRNA expression across B-cell lymphoma models (Figure S20E-F). To validate the relationship between FLI1 and IL-16, we performed a stable knockdown of FLI1 in the ibrutinib-resistant cell line VL51 using shRNA, observing downregulation of the interleukin (Figure 7D). Finally, the ibrutinib-resistant cells were more sensitive than the parental cells to chelerythrine, an antitumor agent that inhibits FLI1 ^42^, and the combination with ibrutinib was beneficial in both parental and resistant cells, with enhanced synergistic effect in the latter (Figure S20G-H). All these data support the role of FLI1 in the resistance to ibrutinib in lymphoma cells.

**Figure 7.**
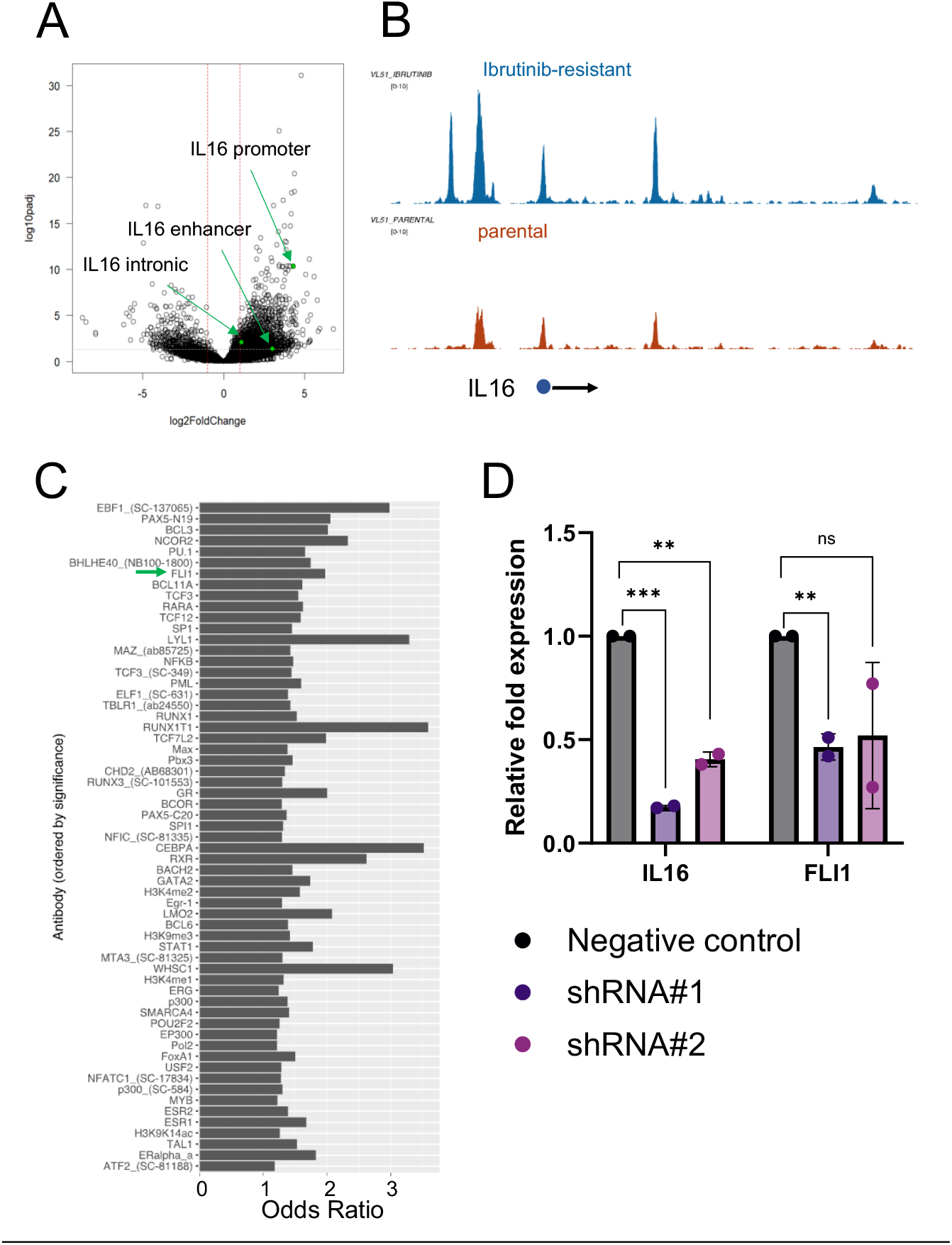
Chromatin remodeling facilitates IL-16 transcription through the FLI1 transcription factor. (A) Volcano plots showing chromatin differential accessibility in the set of consensus peaks based on ATAC-seq data. Dashed lines indicate significance thresholds (*absolute log2 Fold Change > 1, adjusted p-value < 0*.*05*). Highlighted are differentially accessible peaks at the *IL16* locus in the VL51 ibrutinib-resistant cell line compared to its parental counterpart. (B) Chromatin accessibility tracks around the *IL16* locus for VL51 ibrutinib-resistant (top) and parental (bottom). Enrichment is visualized as reads per bins per million base pairs (BPM). The black arrow indicates the transcription start site of the *IL16* gene. (C) Region set enrichment analysis of differentially accessible regions in the VL51 ibrutinib-resistant model, performed using LOLA software, based on publicly available ChIP-seq data (ENCODE, CODEX). Results are ranked by significance (*q*-value). (D) Relative mRNA expression of IL-16 and FLI1 (normalized to GAPDH) upon FLI1 silencing by two different shRNAs in the VL51 ibrutinib-resistant model, compared to the negative shRNA control. * for p<0.05 from a t-test comparing each shRNA vs negative control.

## Discussion

The development and characterization of a novel model of secondary resistance to BTK-targeting agents in splenic MZL led to the identification of the IL-16/CD9 axis as a novel resistance mechanism to targeted agents and chemotherapy in B-cell lymphomas, which can be pharmacologically inhibited.

The development of resistance to BTK inhibitors remains a major therapeutic challenge in patients with B-cell malignancies. We established and characterized an ibrutinib-resistant MZL cell line, VL51-RES, which exhibited a 15-fold increase in IC50 compared to the parental VL51 cells. Resistance was stable even after three weeks in a drug-free medium, suggesting a durable adaptation rather than a transient drug-tolerant phenotype. Furthermore, VL51-RES cells were resistant to covalent and non-covalent BTK inhibitors and BTK degraders.

Genomic profiling of VL51-RES did not reveal mutations in *BTK* or *PLCG2*, two of the most frequently implicated genes in ibrutinib resistance ^1,3,6^. Instead, transcriptomic and epigenetic analyses revealed a profound shift in gene expression, with upregulation of genes involved in cytokine signaling, NF-κB activation, cell proliferation, and PI3K-AKT signaling. Among the most highly upregulated cytokines, IL-16 was prominently increased at both the RNA and protein levels, with concomitant downregulation of its negative regulatory miRNAs, miR-125a and miR-4451 ^29^. Functionally, conditioned media from VL51-RES cells conferred ibrutinib resistance to parental VL51 cells, suggesting a paracrine effect. ELISA identified IL-16 as a key secreted component, and exposure to recombinant IL-16 was sufficient to induce resistance in VL51 parental cells, as well as other CD9-expressing B-cell lymphoma models, including MEC1 (CLL) and REC1 (MCL). These findings suggest that IL-16 secretion by resistant cells may contribute to a protective microenvironment that shields tumor cells from BTK inhibition. Supporting the clinical relevance of this mechanism, serum IL-16 levels were elevated in ibrutinib-resistant CLL patients who, similarly to VL51-RES, lacked *BTK* or *PLCG2* mutations, implicating IL-16 as an alternative route toward resistance.

IL-16 has been linked to inflammatory processes and lymphoid malignancies, though its role in BTK inhibitor resistance remains unexplored. IL-16 expression is primarily restricted to lymphoid tissues, where it functions as a chemoattractant and modulator of T-cell activation. High IL-16 levels have been associated with immune-mediated disorders ^43^, cutaneous T cell lymphomas ^44^, a high disease burden in CLL and multiple myeloma patients ^30,45-47^ and, in multiple myeloma, IL-16 blocking can decrease cell proliferation ^30,46^. Furthermore, in mouse models of colorectal cancer with high lymphocyte infiltration, IL-16 neutralization reduced tumor growth and enhanced antitumor immune populations ^48^. Our data and the literature indicate a complex cytokine-driven resistance network in B-cell malignancies, where IL-16 contributes to cell-intrinsic survival and a broader immunomodulatory microenvironment that can promote resistance to targeted therapies like ibrutinib.

We showed that IL-16 worked via its non-canonical receptor CD9 and not the canonical CD4 expressed in T cells. CD9 is a tetraspanin protein that regulates cell adhesion, migration, and immune signaling, inducing phosphorylation of PI3K and AKT, and mediating its effects on cell migration and immune responses ^49-52^. CD9 is expressed in ABC DLBCL, Waldenstrom’s Macroglobulinemia, and multiple myeloma cells, and its high expression can be associated with inferior outcomes ^30,53,54^, as we also observed using lymphoma bulk and single-cell RNA-Seq datasets.

We showed that blocking IL-16 using a neutralizing antibody or targeting CD9 with a CD9-binding peptide restored ibrutinib sensitivity in resistant cells, indicating that the IL-16/CD9 axis plays a pivotal role in sustaining drug resistance but also providing a potential strategy to tackle the decreased drug activity. Besides giving resistance to BTK inhibitors in MZL, MCL, and CLL models, we found that the IL-16/CD9 signaling also impacted R-CHOP sensitivity in ABC-DLBCL models, reinforcing the broader relevance of this pathway across B-cell malignancies.

Mechanistically, IL-16 stimulation led to NF-κB and PI3K-AKT activation, two pathways that are critical for B-cell survival and proliferation. This was further corroborated by transcriptomic data, which showed upregulation of NF-κB and MYC target genes and repression of oxidative phosphorylation (OxPhos) and interferon signaling. IL-16 also induced CXCL10 secretion, another NF-κB target, further contributing to ibrutinib resistance.

Epigenetic analyses suggested that IL-16 upregulation in resistant cells was driven by increased activity of the transcription factor FLI1, a member of the ETS family. Knockdown of FLI1 reduced IL-16 expression, indicating that chromatin remodeling at the *IL16* gene locus may be a key adaptive mechanism in resistance.

Morphologically, VL51-RES cells were smaller than their parental cells but had increased membrane protrusions and filopodia, which are features commonly associated with increased motility and tumor aggressiveness. In addition to their structural role in invasion, membrane protrusions actively engage in signal transduction pathways, including B-cell receptor signaling ^55^. These observations are consistent with previous reports linking ibrutinib resistance to cytoskeletal remodeling and actin polymerization changes in CLL and MCL ^8,56^. In addition, we observed mitochondrial alterations in resistant cells, including increased size and fragmentation, suggesting enhanced mitochondrial activation. Mitochondrial reprogramming has been increasingly recognized as a hallmark of drug-resistant phenotypes, particularly in hematological malignancies, where it supports metabolic plasticity and survival under therapeutic pressure ^57,58^. Further supporting this link, IL-16 signaling, through its interaction with CD9, induces actin reorganization and promotes cell migration ^59^. Since mitochondria are highly responsive to extracellular signaling events, IL-16 may contribute to mitochondrial adaptations that sustain metabolic flexibility and apoptotic resistance ^60^. Consistently, recombinant IL-16 has been shown to upregulate genes involved in actin polymerization and mitochondrial biogenesis in lymphoma cells ^61^.

Our findings highlight the intricate relationship between cytokine secretion, cytoskeletal remodeling, mitochondrial reprogramming, and therapy resistance in B-cell lymphoma. Targeting these interconnected pathways, which the tumor microenvironment can also sustain ^53^, could offer novel therapeutic strategies to overcome drug resistance and improve treatment efficacy in B-cell malignancies. Indeed, our findings align with emerging evidence implicating inflammatory cytokines in mediating resistance to targeted therapies. Previous studies have demonstrated that CCL3 and CCL4 promote ibrutinib resistance in CLL by creating a pro-survival niche ^62,63^. Other secreted factors, such as IL-6 or heparin-binding epidermal growth factor-like growth factor can sustain resistance to various targeted agents, including BTK and/or BTK inhibitors and CD37 targeting ^9,11,64-66^. Our current results extend these findings by identifying IL-16 as a novel driver of resistance, highlighting the therapeutic potential of targeting cytokine-driven resistance mechanisms.

Clinically, our findings suggest that IL-16 may serve as a biomarker to identify patients at risk of developing resistance to BTK inhibitors, particularly those who lack BTK or PLCG2 mutations. Paired with circulating tumour DNA analysis ^67^, measuring serum IL-16 levels in patients undergoing BTK inhibitor therapy could help predict resistance early, potentially guiding future treatment decisions. Furthermore, our results provide a strong rationale for developing IL-16 or CD9-targeting therapies, either as monotherapies or combined with BTK inhibitors or chemoimmunotherapy. Future directions should focus on prospectively validating these findings in patient samples and assessing whether IL-16 inhibition can enhance BTK inhibitor efficacy in clinical settings. Given the role of IL-16 in modulating the tumour microenvironment, additional studies should explore its impact on immune cell interactions and whether IL-16 blockade can enhance responses to immune-based therapies.

In conclusion, our study identifies IL-16/CD9 as a novel resistance mechanism to BTK inhibitors and R-CHOP in B-cell lymphomas. Targeting this axis may provide a promising strategy to overcome resistance and open new therapeutic possibilities to be explored in the clinical setting.

## Author contribution

AJA performed experiments, analyzed and interpret data, performed data mining, prepared the figures and cowrote the manuscript; FG performed ATAC-seq data analyses and interpreted data; LC, LTB and XY performed data mining; EC, SN, GS, FF, CT, FS, performed experiments; AZ, FR, RBP, GS, VG performed flow-cytometry analyses; AB performed variant calling, AR performed genomics experiments; MCM, ME performed methylation profiling experiments and data mining, SJ performed ELISA Luminex analyses; AS provided advice; YBD and FS provided reagents and advice; JRB and QPH collected and characterized tumor samples; EZ and DR codesigned research, edited the manuscript; FB designed research, interpreted data, and cowrote the manuscript; all authors approved the final manuscript.

