## Supplementary figures and tables for "IL-16 production is a mechanism of resistance to BTK inhibitors and R-CHOP in lymphomas"

### Supplementary tables

**Table S1**. Panel of antibodies used in flow cytometry experiments.

| Protein | fluorochrome | company | # catalog |
|---|---|---|---|
| CD4 | PE-Cy7 | BD Bioscience | 348809 |
| CD9 | V450 | BD Bioscience | 561326 |
| CXCR3 | APC | BD Bioscience | 561732 |
| IL-16 | Alexa350 | R&D Systems | IC3161U |

**Table S2**. Panel of proteins tested in the immunoblotting experiments.

| source | protein | company | # catalog |
|---|---|---|---|
| rabbit polyclonal | α-AKT | Cell Signaling | 9272 |
| rabbit polyclonal | α-p(S473) AKT | Cell Signaling | 4060 |
| mouse monoclonal | α-vinculin | Merk | V9131 |
| Rabbit monoclonal | RELA NF-κB/p65 (C22B4) | Cell Signaling | 4764 |
| Rabbit monoclonal | RELB (C1E4) | Cell Signaling | 4922 |
| Rabbit monoclonal | NF-κB1 p105/p50 (D4P4D) | Cell Signaling | 13586 |
| Rabbit polyclonal | NF-κB2 p100/p52 | Cell Signaling | 4882 |

#### Supplementary Figures

**Figure S1**. **Profiles of drug sensitivity differ between parental and resistant cell lines**. Acquired resistance was tested by MTT assay (72 hours) in parental (black) and resistant (red dashed) VL51 cells. Drug sensitivity was evaluated in resistant and parental cells for the BTK inhibitors ibrutinib (A), zanubrutinib (B), acalabrutinib (C), pirtobrutinib (D), and the BTK degrader NX-5948 (E). Error bars correspond to the standard deviation of the mean; μM for micromolar. Data was derived from at least three independent experiments. P values from a Z-test; statistically significant for p< 0.05.

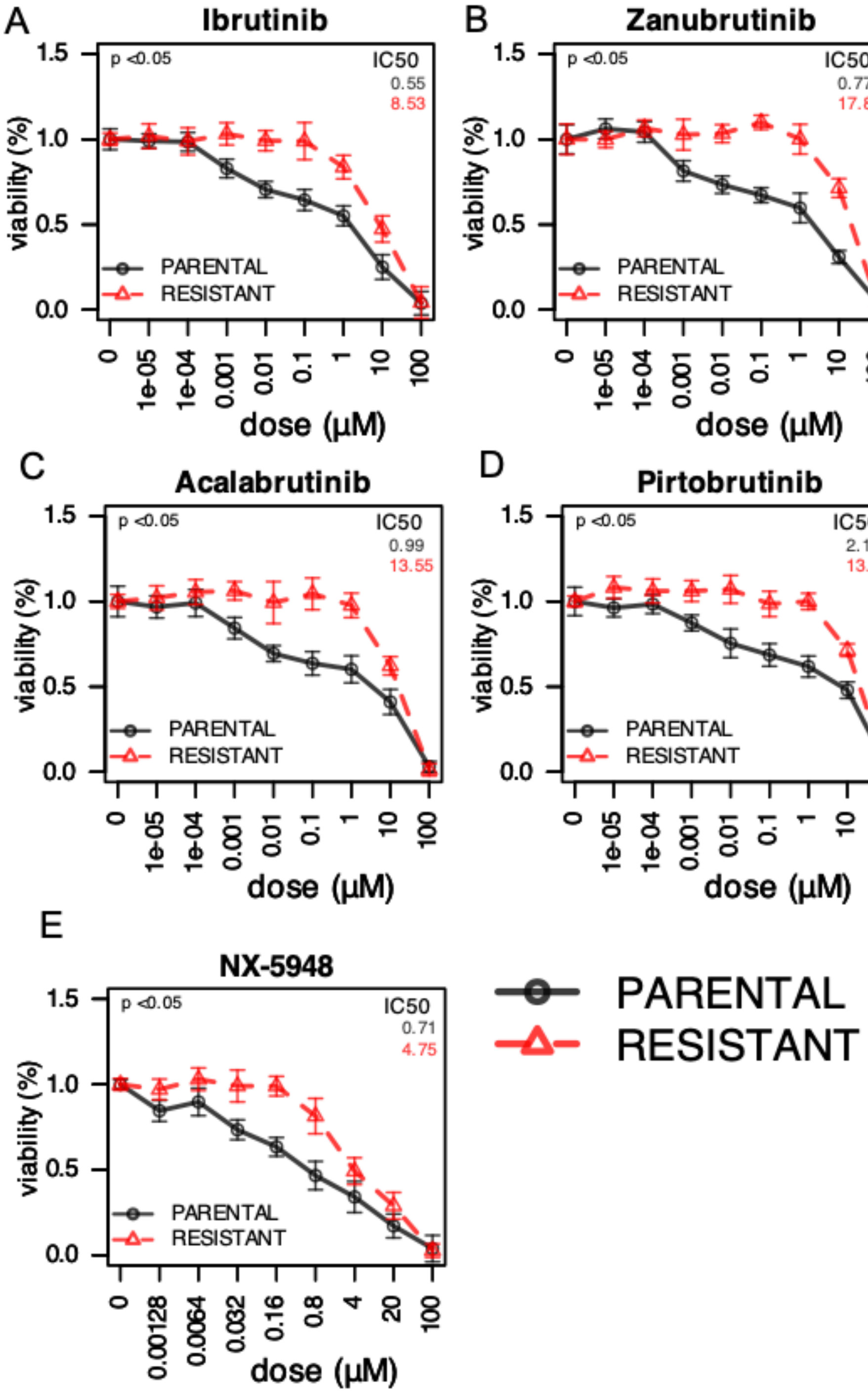

**Figure S2**. **Multi-drug resistance assessment.** Multi-drug resistance (MDR) phenotype was ruled out by the expression of MDR1 (blue) and MDR2 (orange) genes by real-time PCR in VL51 parental and ibrutinib-resistant cells (A), and by the sensitivity to the chemotherapy agent and MDR substrate vincristine (B). The barplot in (A) represents the mean of RQ values (normalized expression) calculated by the DDCt method. Data is derived from two independent experiments; error bars represent the standard deviation of the mean. The graph in (B) represents the drug-response curve to vincristine in VL51 parental cells (black) and ibrutinib-resistant cells in dashed red. Error bars correspond to the standard deviation of the mean; µM for micromolar. Data was derived from at least three independent experiments. P values from a Z-test are statistically significant for $p < 0.05$.

A

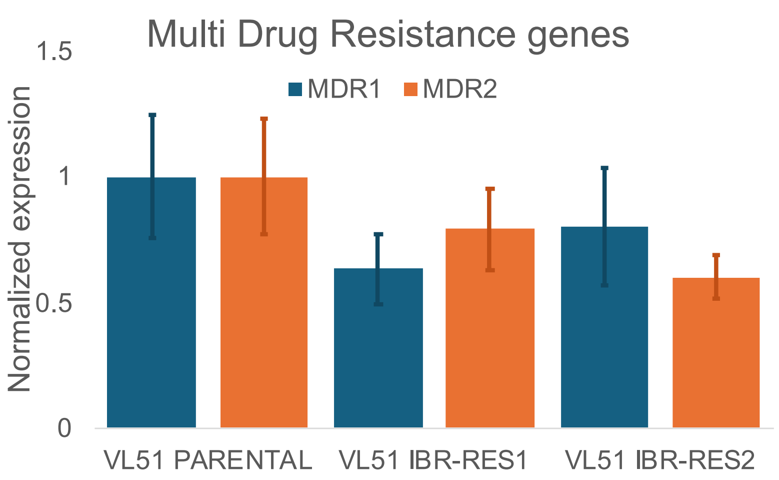


B

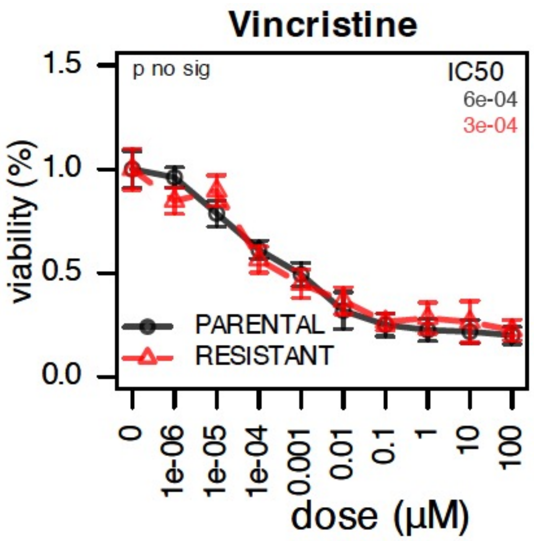

**Figure S3. Morphometry of VL51 parental (black) and ibrutinib-resistant VL51 cells (blue) by electronic microscopy.** Boxplots represent the quantification of cell size (A top) and membrane protrusion (A bottom), and mitochondrial ultrastructural analyses: area (B left), perimeter (B left-center), major axis length (B right-center), eccentricity (B right). Data was derived from two independent experiments. P values from a t-test are statistically significant for p< 0.05.

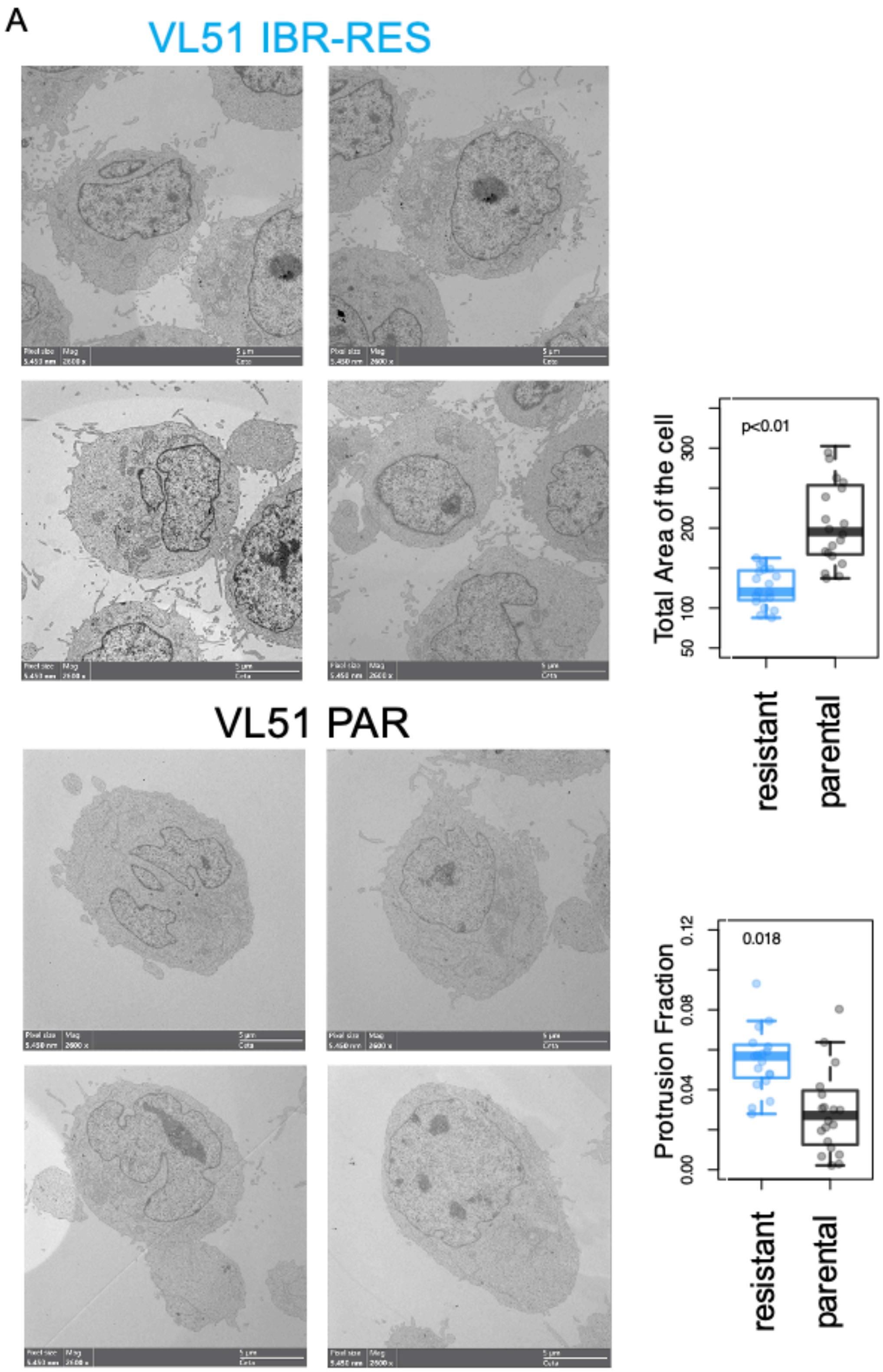

B

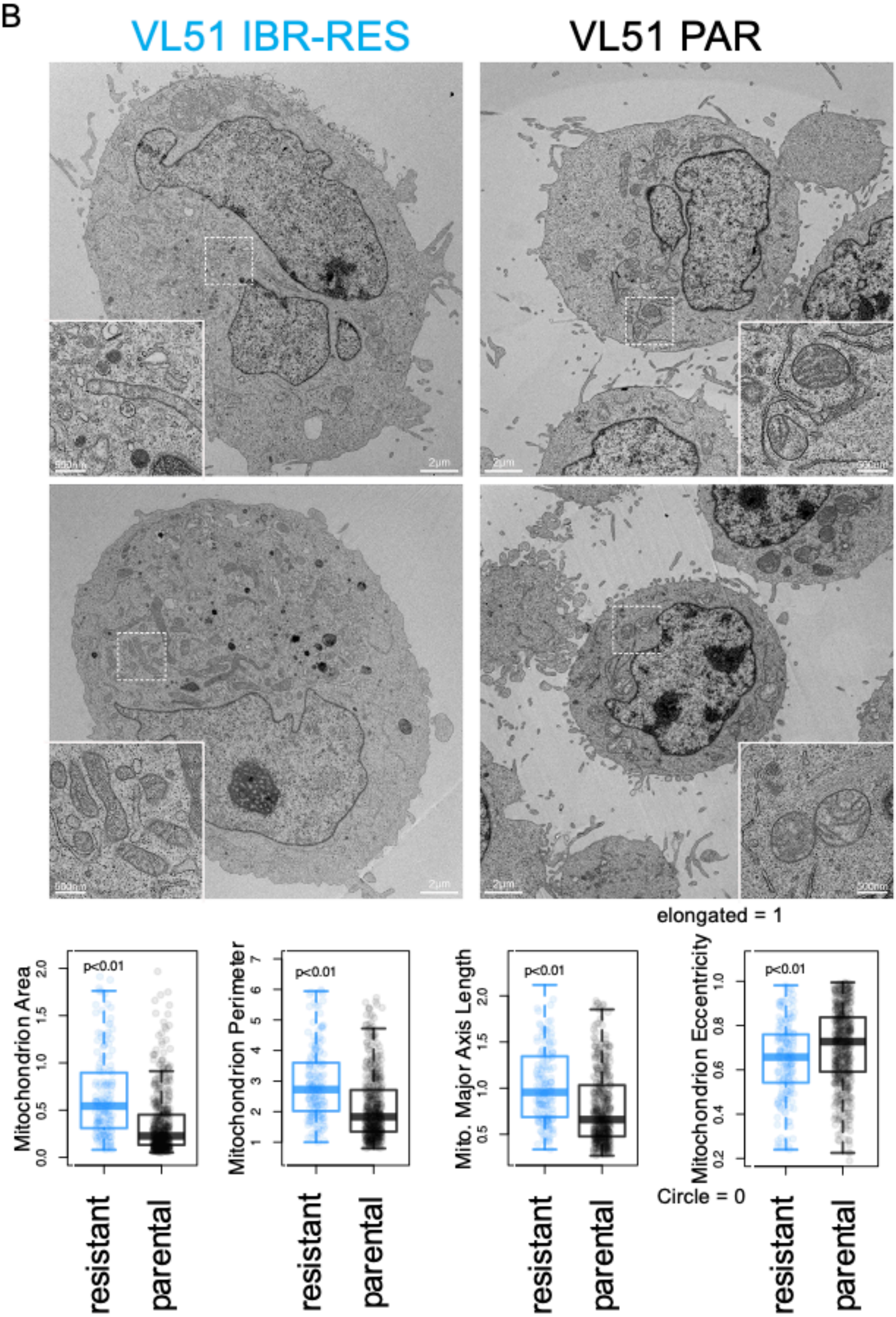

VL51 IBR-RES
VL51 PAR
2µm
500nm
elongated = 1
Circle = 0
Mitochondrion Area
Mitochondrion Perimeter
Mito. Major Axis Length
Mitochondrion Eccentricity
p<0.01
resistant
parental

**Figure S4**. **Profiles of drug sensitivity differ between parental and resistant cells**. The drug response was tested by MTT assay (72h) in parental (black) and resistant (red dashed) VL51 cells. Drug sensitivity was evaluated in resistant and parental cells for the PI3K inhibitors idelalisib (A), umbralisib (B), copanlisib (C), and duvelisib (D). Error bars correspond to the standard deviation of the mean; μM for micromolar. Data was derived from at least three independent experiments. P values from a Z-test are statistically significant for $p < 0.05$.

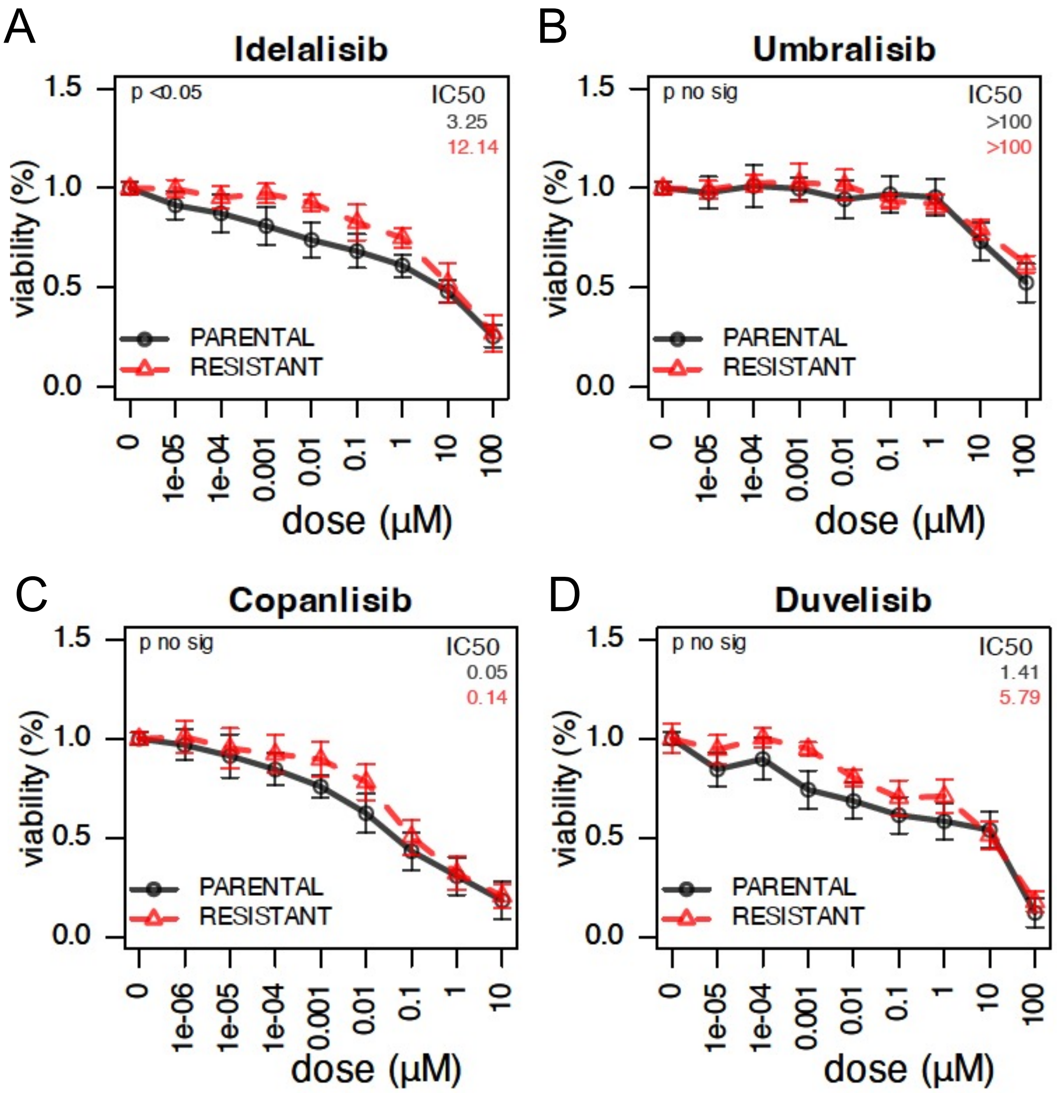

**Figure S5. Conditioned medium from VL51 resistant cells gives resistance to ibrutinib in parental cells.** Parental cells were cultured for 48 hours in the resistant-conditioned medium (from 24 hours of culture in resistant cells), followed by exposure to increasing concentrations of ibrutinib for 72 hours. The plot shows the ibrutinib response in parental (black), parental cultured in conditioned medium from resistant cells (black dashed), and resistant (red) VL51 cells. Error bars correspond to the standard deviation of the mean; μM for micromolar. Data was derived from at least three independent experiments. P values from a Z-test, statistically significant for $p < 0.05$.

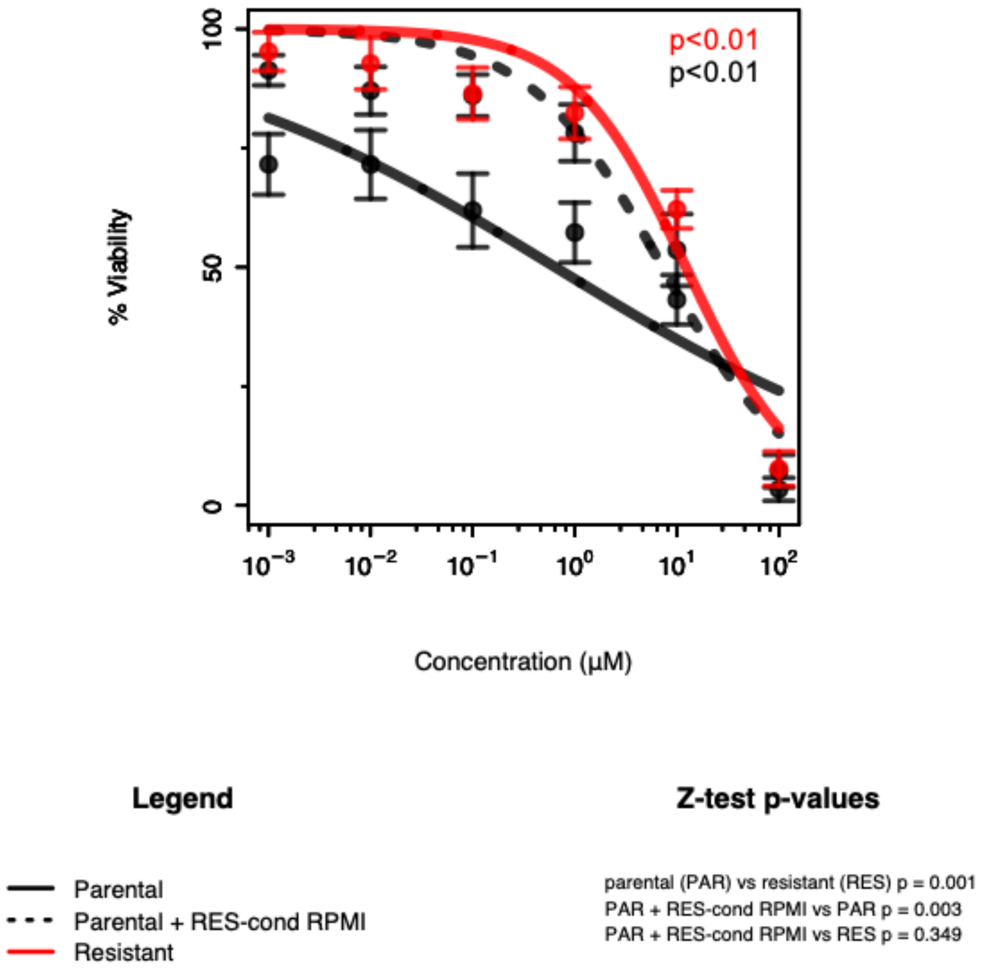

**Figure S6. Secreted levels of cytokines were measured using the Proteome Profiler Human Cytokine Array Kit (R&D Systems) following the manufacturer's protocol.** The barplot shows the normalized expression for resistant and parental conditioned medium from the cytokine array membranes. Error bars represent the standard deviation of the mean. Data was derived from two independent experiments. P values from the Welch Two Sample test, * for statistically significant ($p < 0.05$).

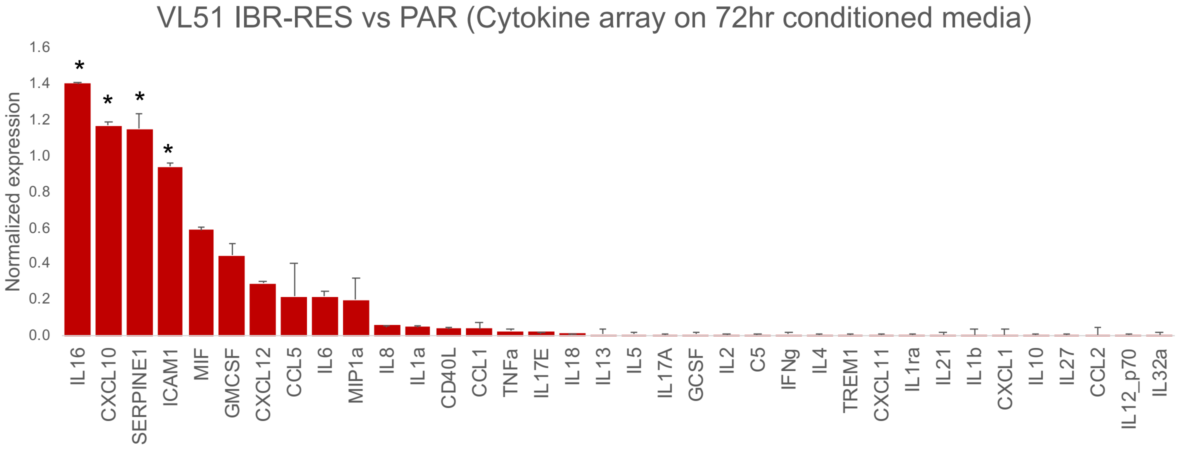

**Figure S7. Expression levels of intracellular IL-16 were measured by flow cytometry**. (A) Scatter plot on IL-16 expression (Alexa350, log10) in VL51 parental (left) and ibrutinib-resistant (right). (B) Density plots show the median MFI values of three replicates: negative control (dashed black), parental (grey), and resistant (red). P-value from the Welch Two Sample test performed on MFI values of the three experiments.

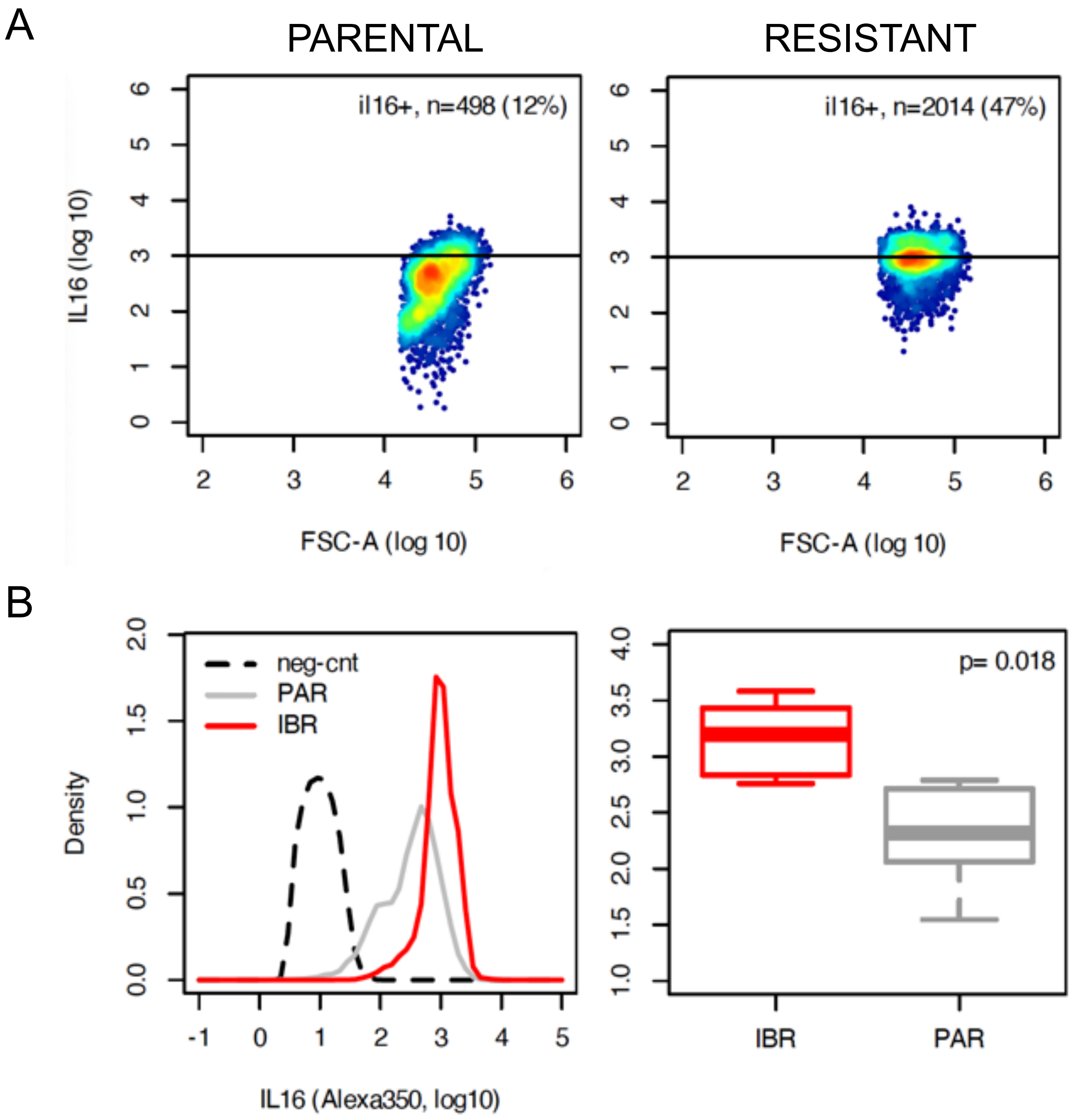

**Figure S8. NFkB1, NFkB2, and pAKT/AKT protein levels in parental and resistant VL51 cells.** (A) Immunoblotting was performed to measure protein levels in parental and resistant clones for NFkB1, NFkB2, and pAKT/AKT in two independent experiments. Vinculin was used as a loading control. (B) Protein quantification was done, normalizing first to vinculin and then to parental levels. The barplot represent the mean of the two independent experiments, parental in grey, resistant in orange. Error bars represent the standard deviation of the mean. Values at the end of the bars represent the p-value from a Welch Two Sample test.

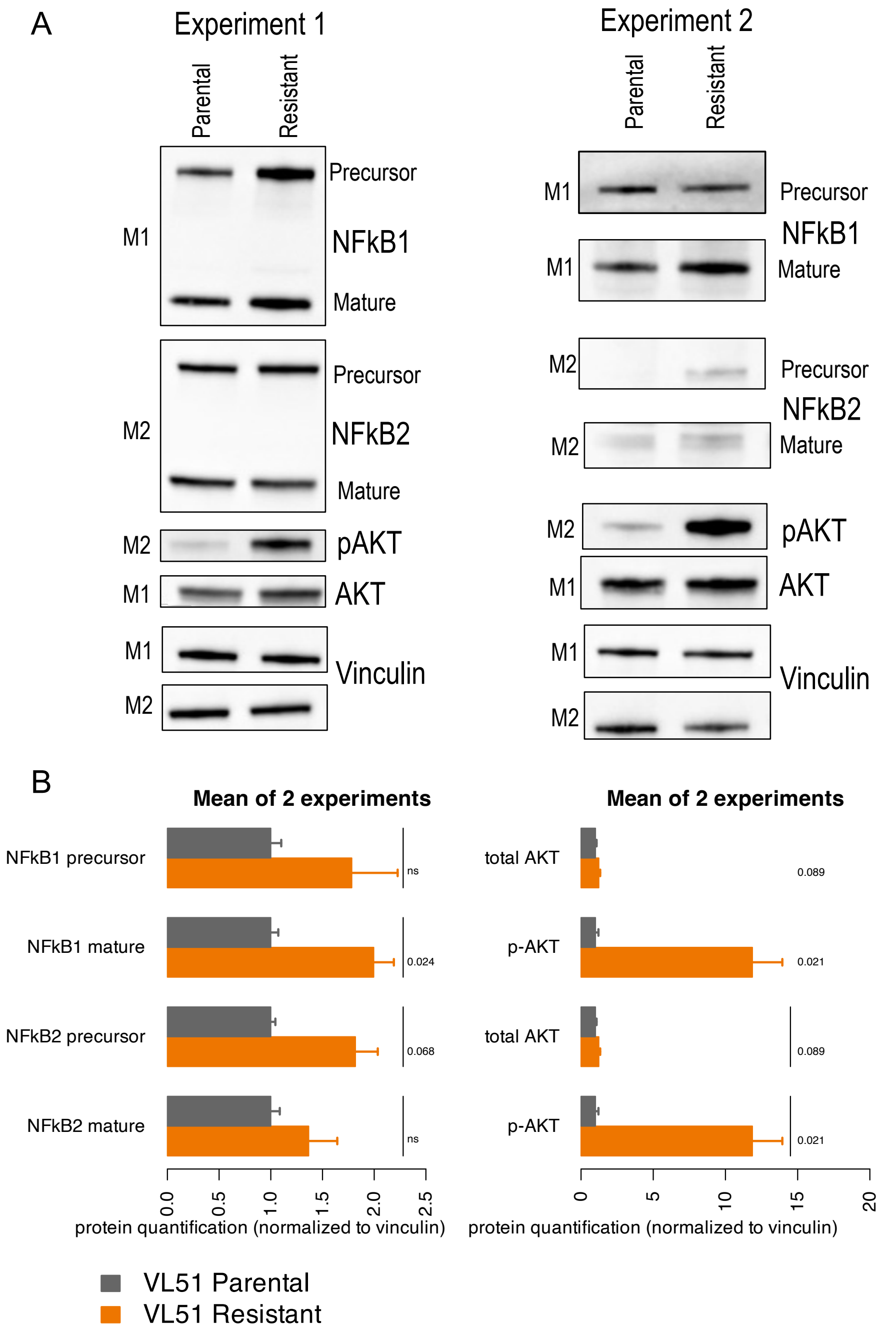

**Figure S9. Expression levels of surface CD4 (A) and CD9 (B) were measured by flow cytometry**. Density plots show the median MFI values of at least two independent experiments. (A) Surface CD4 expression in VL51 parental and ibrutinib-resistant cells: negative control (dashed black), parental (grey), resistant (dashed red). (B) Surface CD9 expression in VL51 parental (grey) and ibrutinib-resistant (two biological replicates in red and orange) cells, and Karpas1718 parental (green) and idelalisib-resistant (blue) cells, negative control in black. Karpas1718 cells were included as a CD9-negative control. (C) Surface CD9 expression in VL51 parental (grey) and ibrutinib-resistant (dashed red) cells, negative control in black.

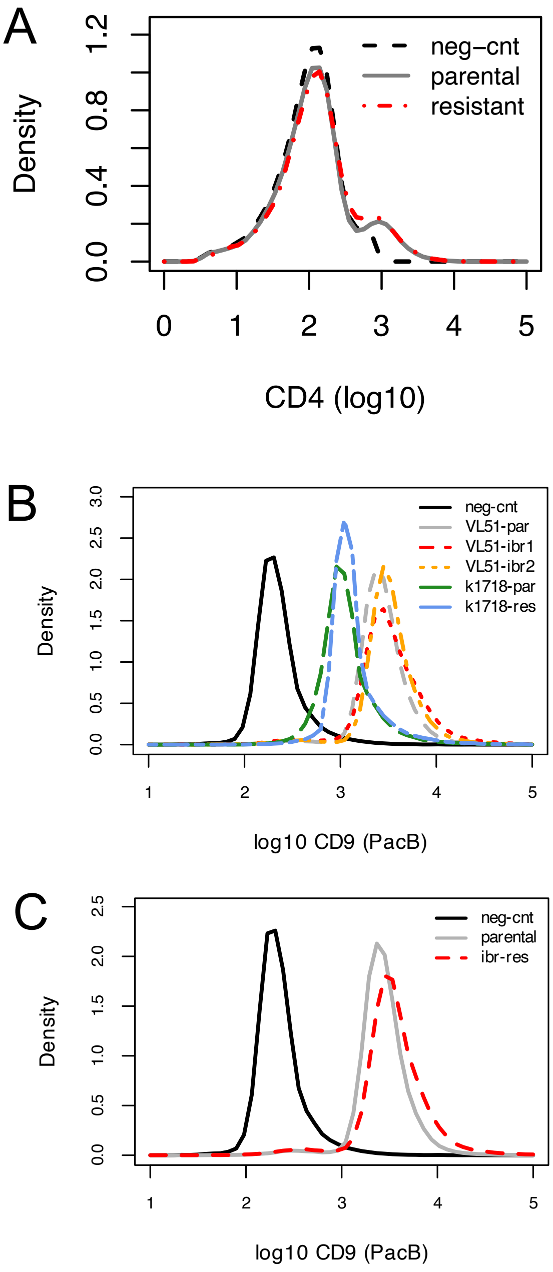

**Figure S10. RNA expression levels of IL-16 and CD9 by RNA-sequencing.** The barplot represents the normalized expression (Z-value) of IL-16 (orange) and CD9 (green) across the B-cell lymphoma cell lines included in the experiments. SMZL for Splenic marginal zone lymphoma, CLL for Chronic lymphocytic leukemia, MCL for Mantle cell lymphoma, and ABC DLBCL for Activated B-cell like diffuse large B-cell lymphoma.

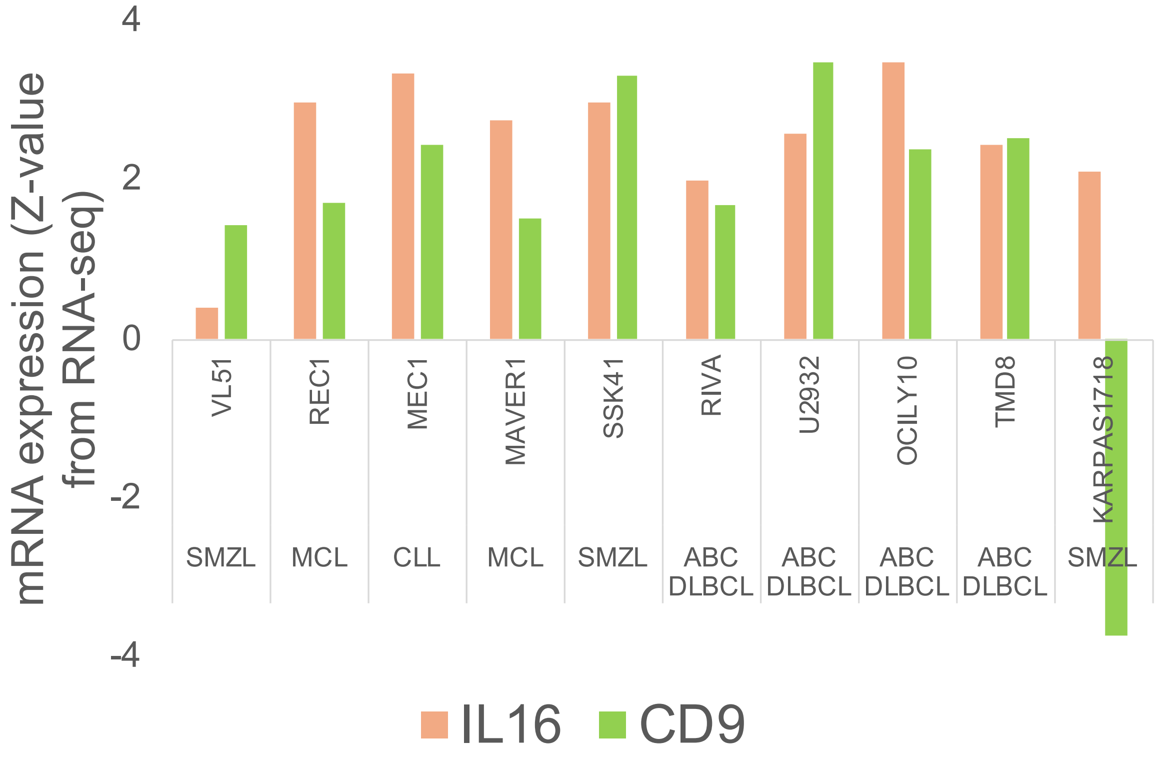

**Figure S11. Gene set enrichment analyses performed on RNA-sequencing data comparing ibrutinib-resistant (red) versus parental (blue) VL51 cells**. Gene expression profile of ibrutinib-resistant showed enrichment of the signature of genes up-regulated in VL51 parental cells upon stimulation with human recombinant IL-16.

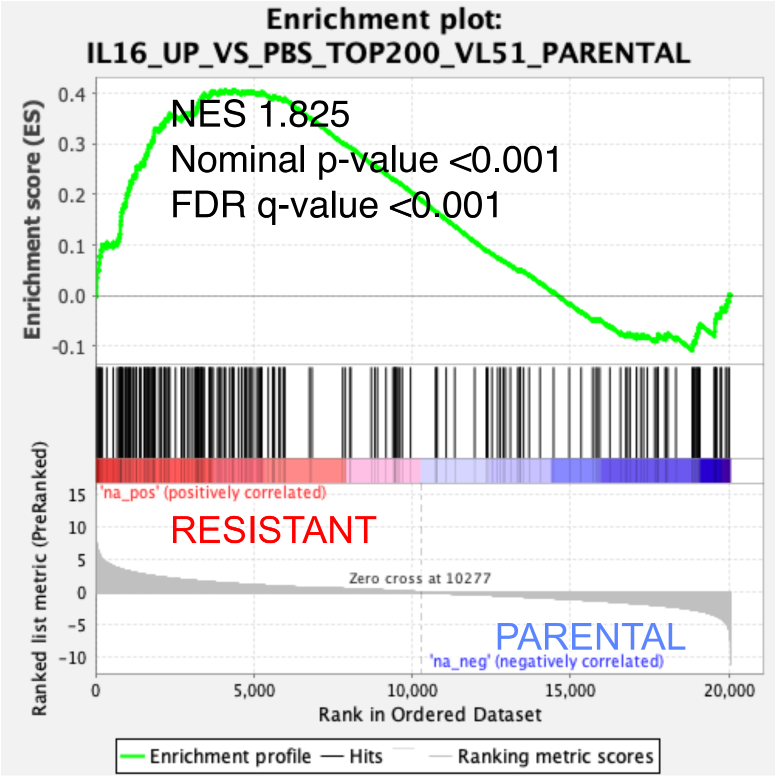

**Figure S12. Gene expression profile of VL51 parental cells upon stimulation with human recombinant IL-16 partially induced the ibrutinib-resistant gene expression profiles.** Venn diagram on the overlap of up-regulated genes in ibrutinib-resistant (IBR-RES_UP, blue), down-regulated genes in ibrutinib-resistant (IBR-RES_DN, green), up-regulated (IL-16_UP, red), and down-regulated (IL16_DN, yellow) genes by recombinant IL-16 in VL51 parental cells. The bottom tables show the commonly up-regulated (red) and down-regulated genes in ibrutinib-resistant cells and by recombinant IL-16 in parental cells.

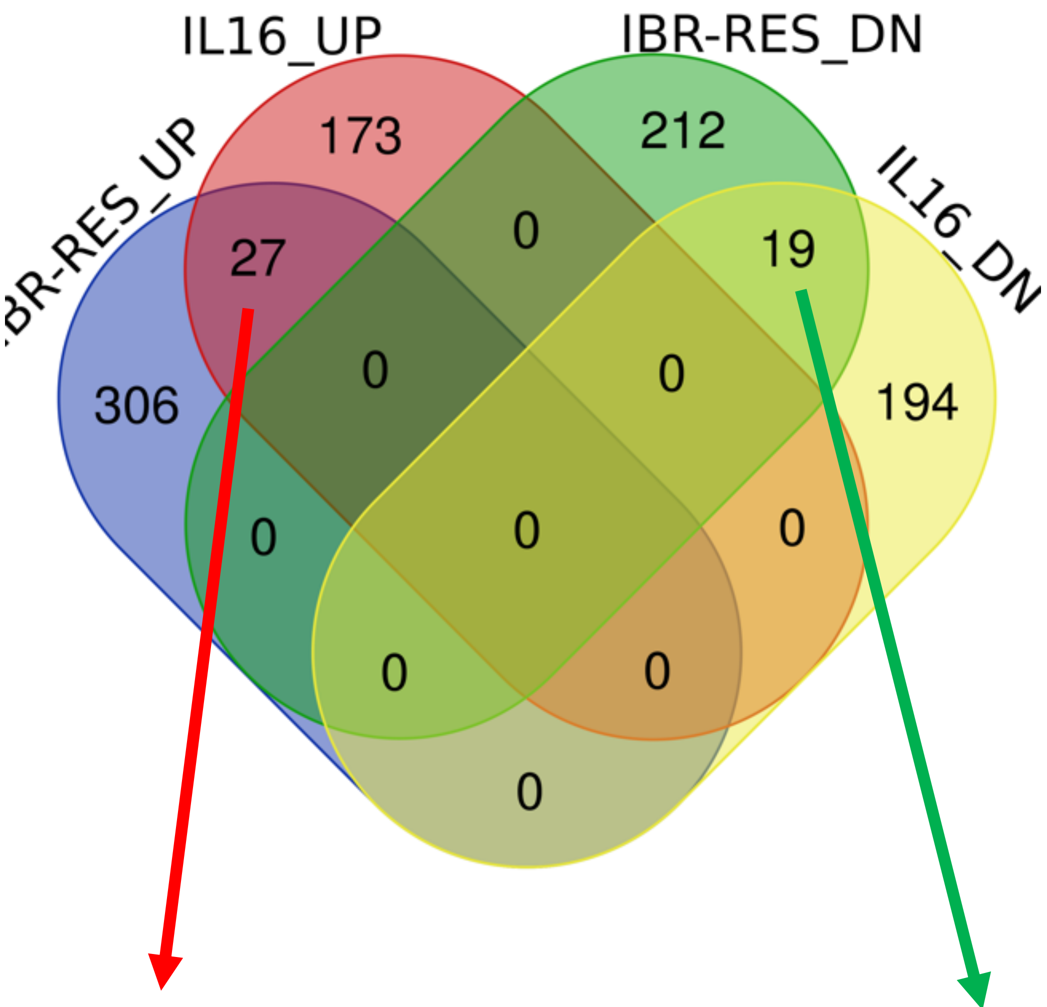


**common UP**

| | | |
|---|---|---|
| PDE1C | LYAR | SLCO4A1 |
| **IGF1** | **BCL2A1** | RRP1B |
| DMBT1 | NOP16 | CDK5R1 |
| SLC2A5 | CD3EAP | POLR3G |
| MYOF | ZFP36L1 | **EGR1** |
| **NR4A1** | PTPN6 | CLN6 |
| **EGR2** | PPP1R16B | CCDC86 |
| **HDAC9** | **TNFRSF13B** | |

**common DN**

| | | |
|---|---|---|
| **TNFAIP2** | CRYZ | **RNF19B** |
| SLC38A2 | **CLIC2** | REPS2 |
| SLAMF6 | **YPEL5** | SLC22A23 |
| **PDCD4** | KCNQ1 | TBC1D9 |
| NECAP2 | CAPG | **METTL7A** |
| EPSTI1 | **DEPTOR** | **ANXA1** |
| OGFRL1 | | |

**Figure S13.** Gene set enrichment analyses performed on RNA-sequencing data: (A) ibrutinib-resistant (red) versus parental (blue) VL51 cells, (B) recombinant IL-16 (red) vs PBS (blue) in VL51 parental, (C) heatmap on genes involved in negative regulation of actin polymerization upon IL-16 stimulation in VL51 parental cells. NES for normalized enrichment score, p for nominal p-value, q for false discovery range.

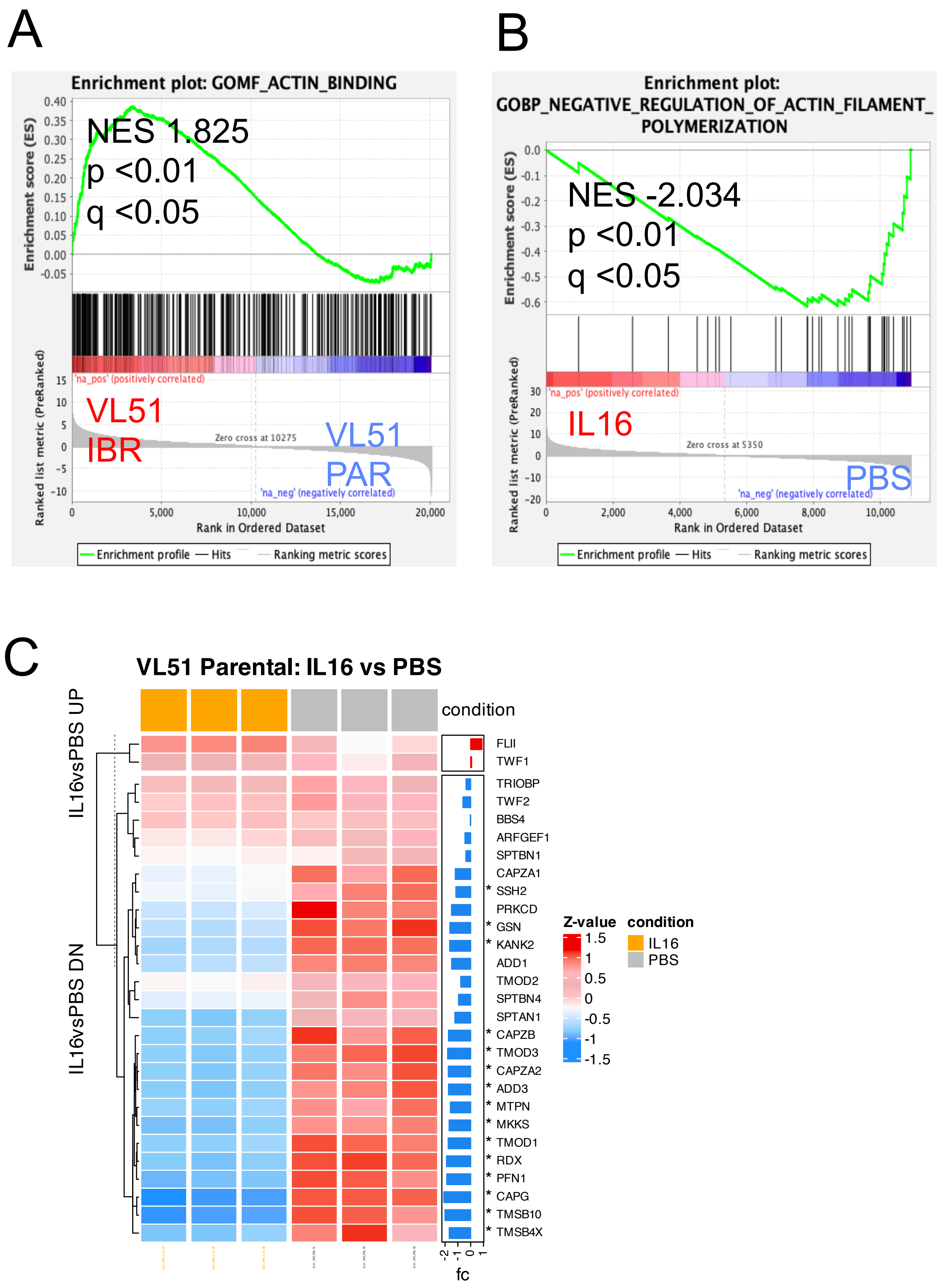

**Figure S14.** Gene set enrichment analyses performed on RNA-sequencing data comparing recombinant IL-16 (red) vs PBS (blue) in VL51 parental: (A) mitochondrial fusion (Gene Ontology biological process signature, (B) heatmap on genes involved in mitochondria biogenesis/activation upon IL-16 stimulation in VL51 parental cells, (C) apoptosis signature (KEGG database). NES for normalized enrichment score, p for nominal p-value, q for false discovery range.

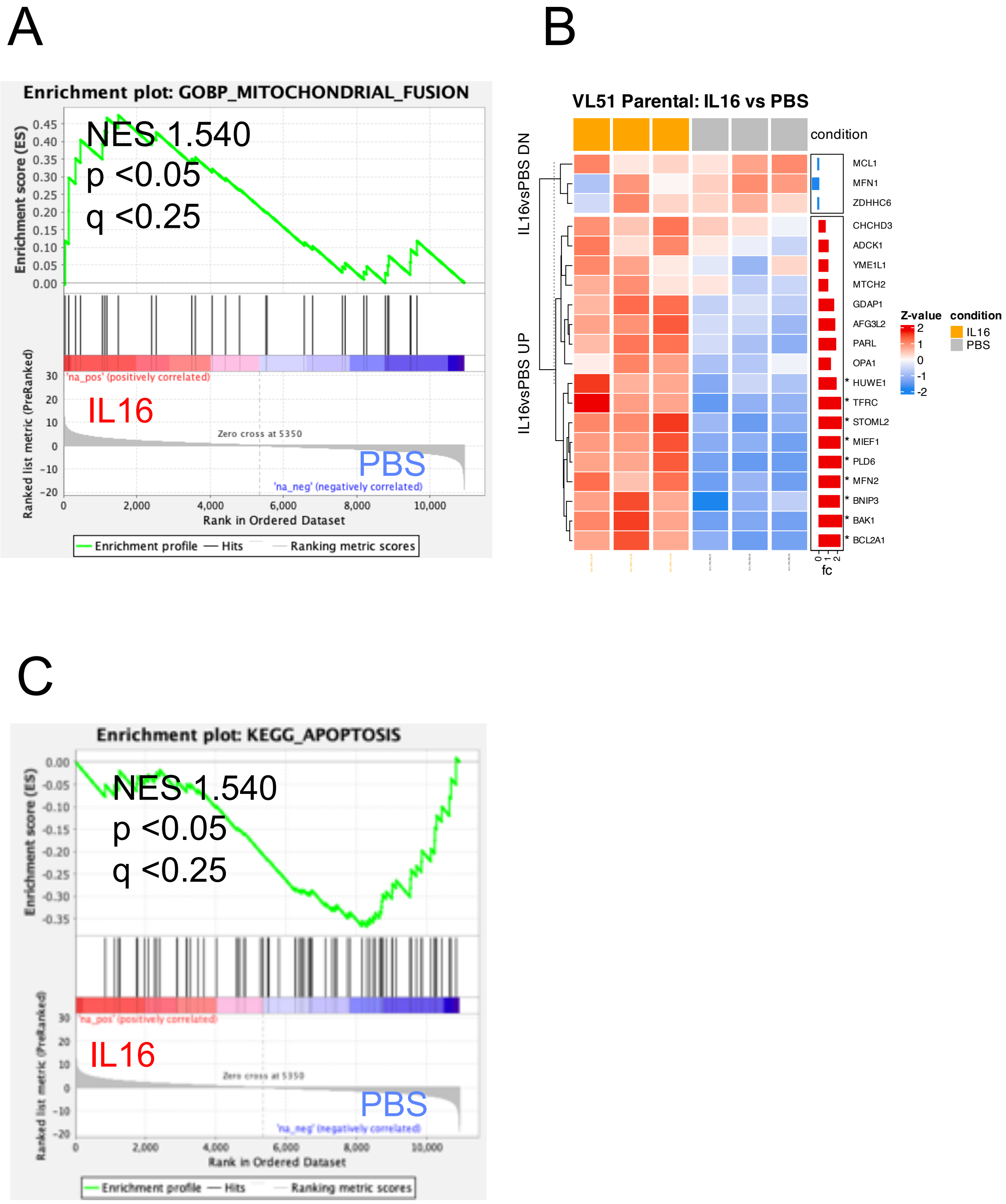

**Figure S15.** Immunoblotting was performed to measure protein levels in parental and resistant clones for NFkB1, NFkB2, RelA and RelB in VL51 parental cells upon human recombinant IL-16 (10ng/mL in yellow, 50ng/mL in orange) or PBS (blue), and in the VL51 ibrutinib-resistant (red) cells. Two independent experiments were performed (A, B). Vinculin was used as a loading control. The barplot on the left show the protein quantification normalized to vinculin.

A Experiment 1

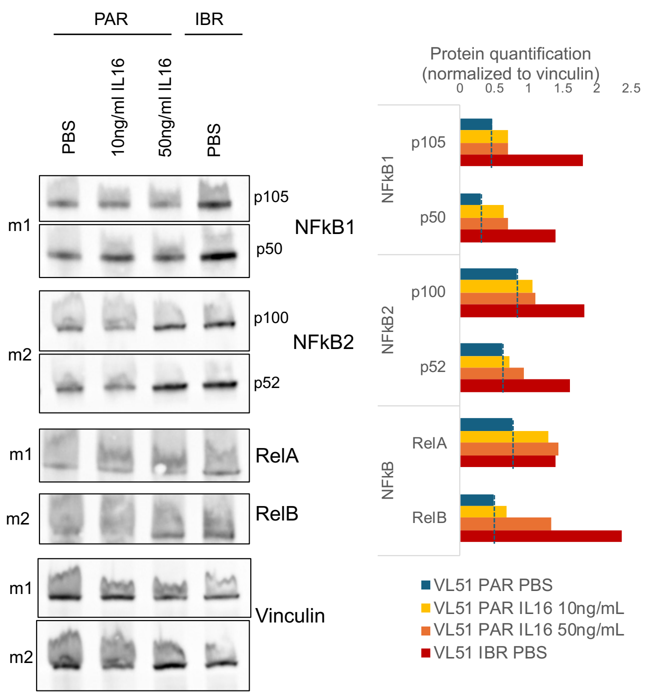


B Experiment 2

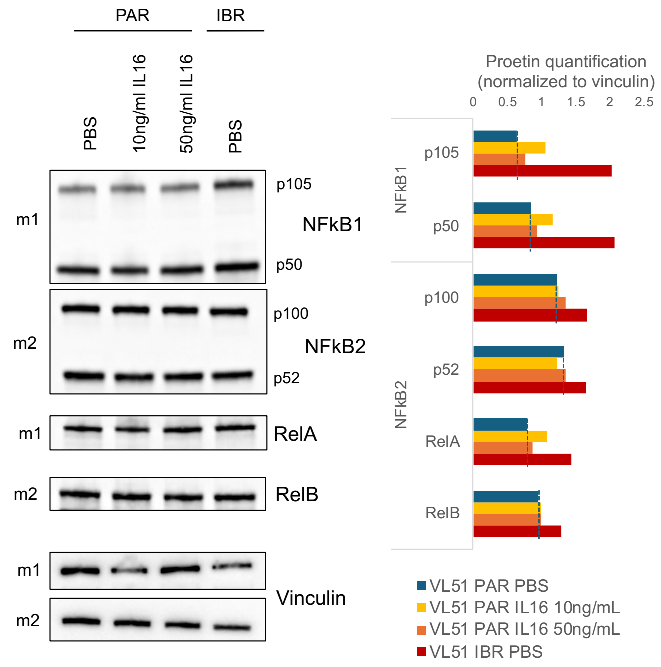

**Figure S16**. (A) Expression levels of surface CXCR3 was measured by flow cytometry. Density plot shows the median MFI values of a representative of at least two independent experiments: VL51 parental (two biological replicates in black and grey) and ibrutinib-resistant (two biological replicates in red and blue) cells. (B) The barplot represents the mean of RQ values (normalized expression) calculated by the DDCt method on the CXCL10 expression upon recombinant IL-16 stimulation (10ng/mL in yellow, 50ng/mL in orange), or PBS (control, grey) for 4 or 24 hours in VL51 parental cells. Data derived from two independent experiments; error bars represent the standard deviation of the mean. (C) ibrutinib-resistant (red, no stimulation) or parental VL51 cells, stimulated with recombinant IL-16 (orange), CXCL10 (yellow), or PBS (control, grey), were exposed to increasing concentrations of ibrutinib for 72 hours. Error bars correspond to the standard deviation of the mean; μM for micromolar. Data derived from at least three independent experiments. P values from a Z-test were statistically significant for p< 0.05.

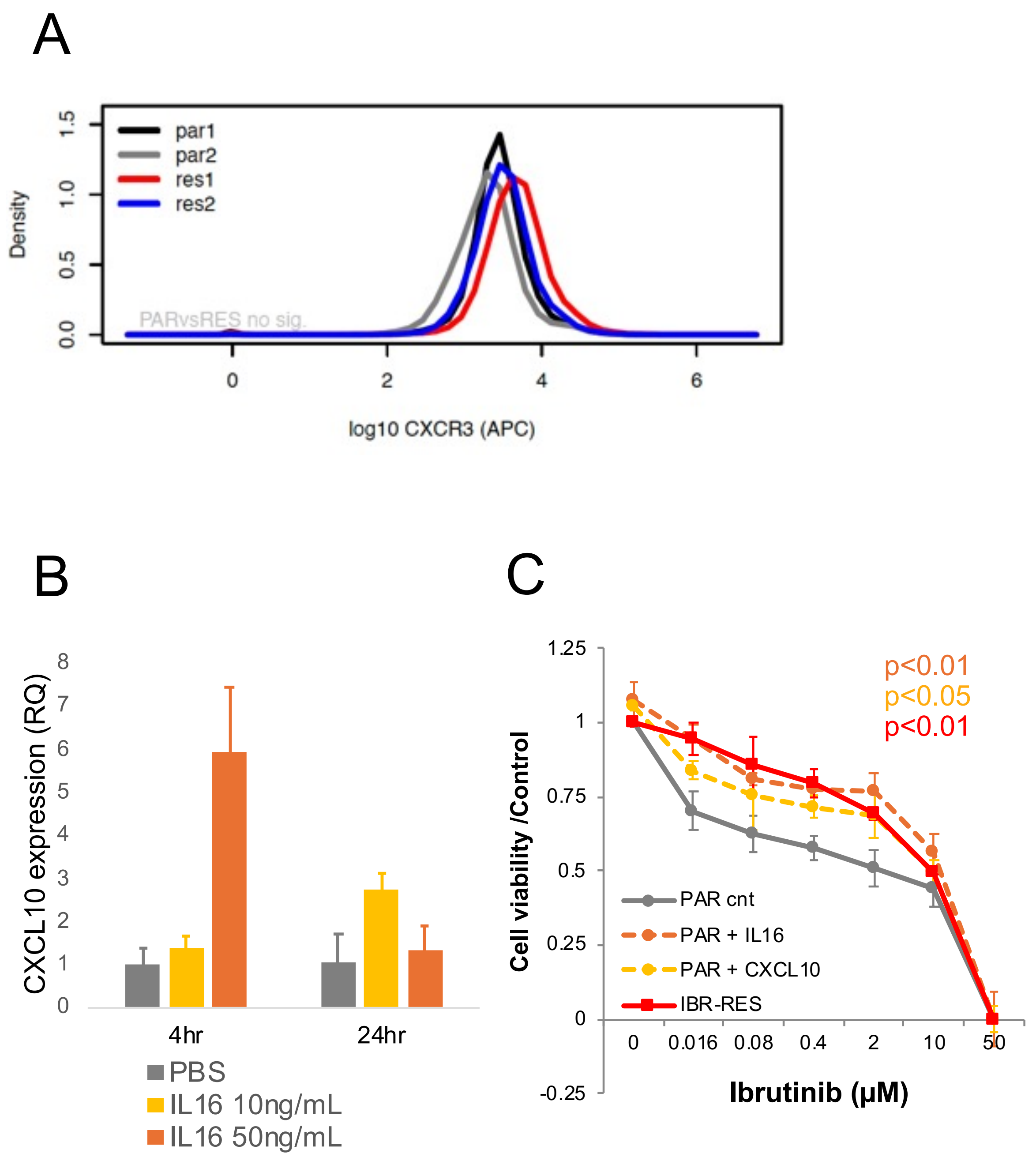

**Figure S17**. Expression levels of IL-16 and CD9 correlated with resistance to ibrutinib. IL-16 (A) and CD9 (B), but not CXCL10 (C), expression levels are inversely correlated with ibrutinib sensitivity. Expression and sensitivity data were analyzed from a previous publication of our group in a panel of 34 B-cell lymphoma cell lines [1]. Cell lines were split into two groups based on whether they had higher or lower values than the median expression of the corresponding gene. Mean of ibrutinib IC50 was calculated for these two groups and compared by t-test.

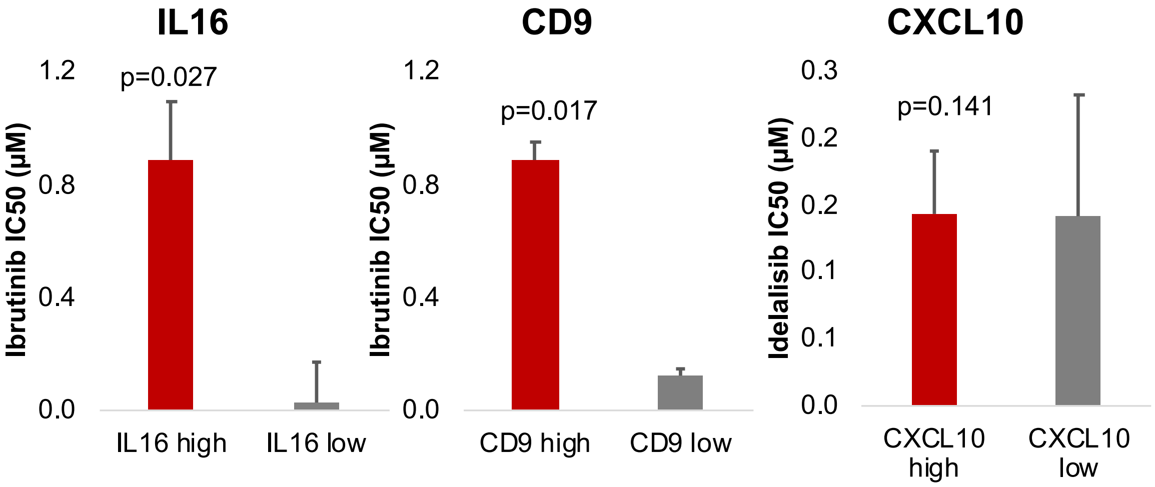

**Figure S18**. Factors associated with resistance to ibrutinib/RCHOP in cell lines are expressed in clinical specimens. (A) Expression levels of genes related to ibrutinib/RCHOP resistance were studied across different subtypes of B cell lymphoma (A) n=80 [2]. B-CLL: chronic lymphocytic leukemia, B. FL: follicular lymphoma, LBNH: non-specified non-Hodgkin B cell lymphoma, MALT: MZL of the mucosa-associated tissue, MCL: Mantle cell lymphoma, NMZL: nodal marginal zone lymphoma, SMZL: splenic marginal zone lymphoma. Red is for highly expressed genes in B cells (CD79A), blue is for genes not expressed in B cells (IGFBP1), and black is for the gene of interest. (B-C) Expression levels of genes related to ibrutinib/RCHOP resistance were studied by single-cell sequencing along different subtypes of non-tumoral immune cells and in malignant cells from the two classic subtypes of diffuse large B-cell lymphoma (DLBCL). ABC: activated B-cell-like DLBCL, GCB: germinal center B-cell-like DLBCL [3].

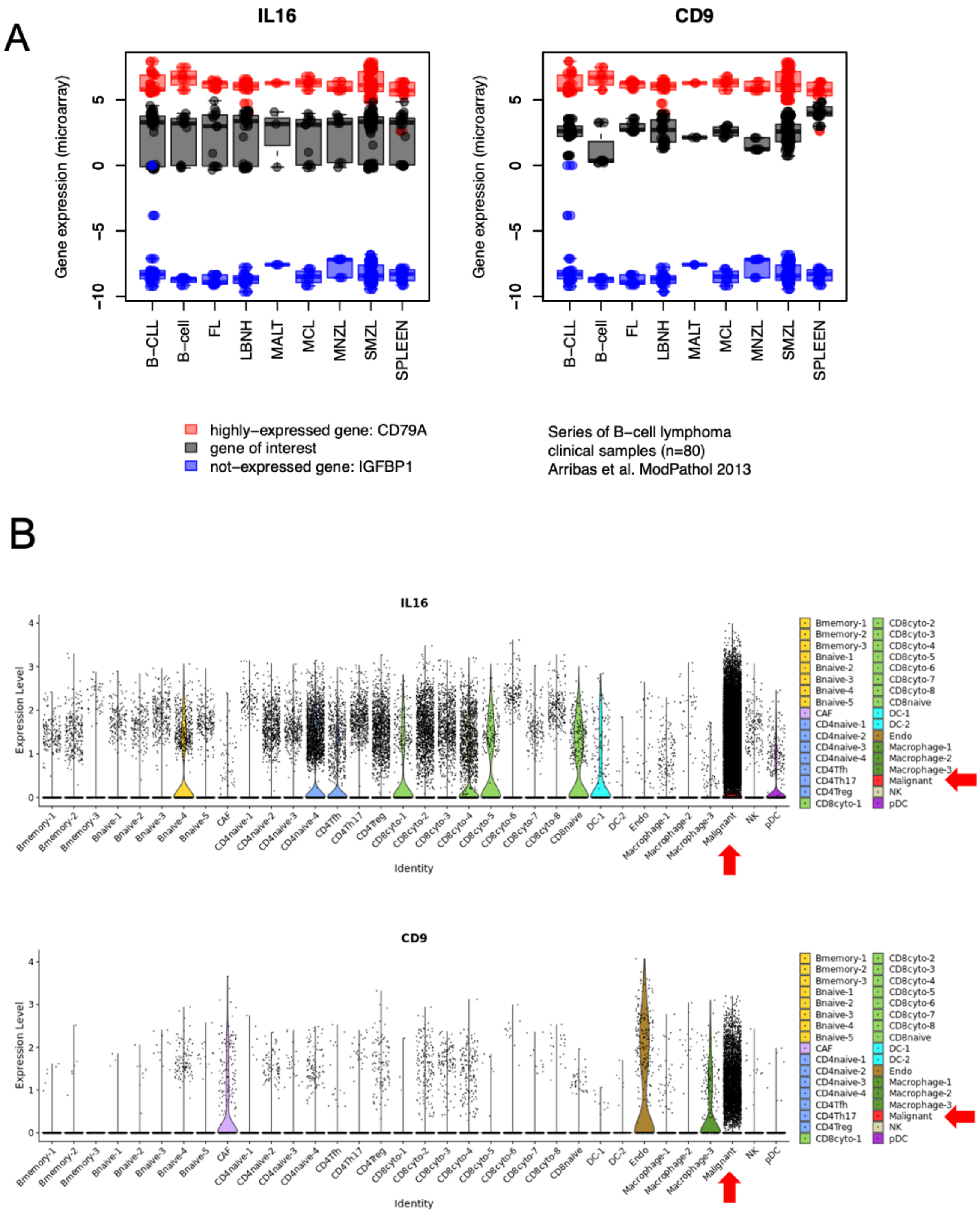

C

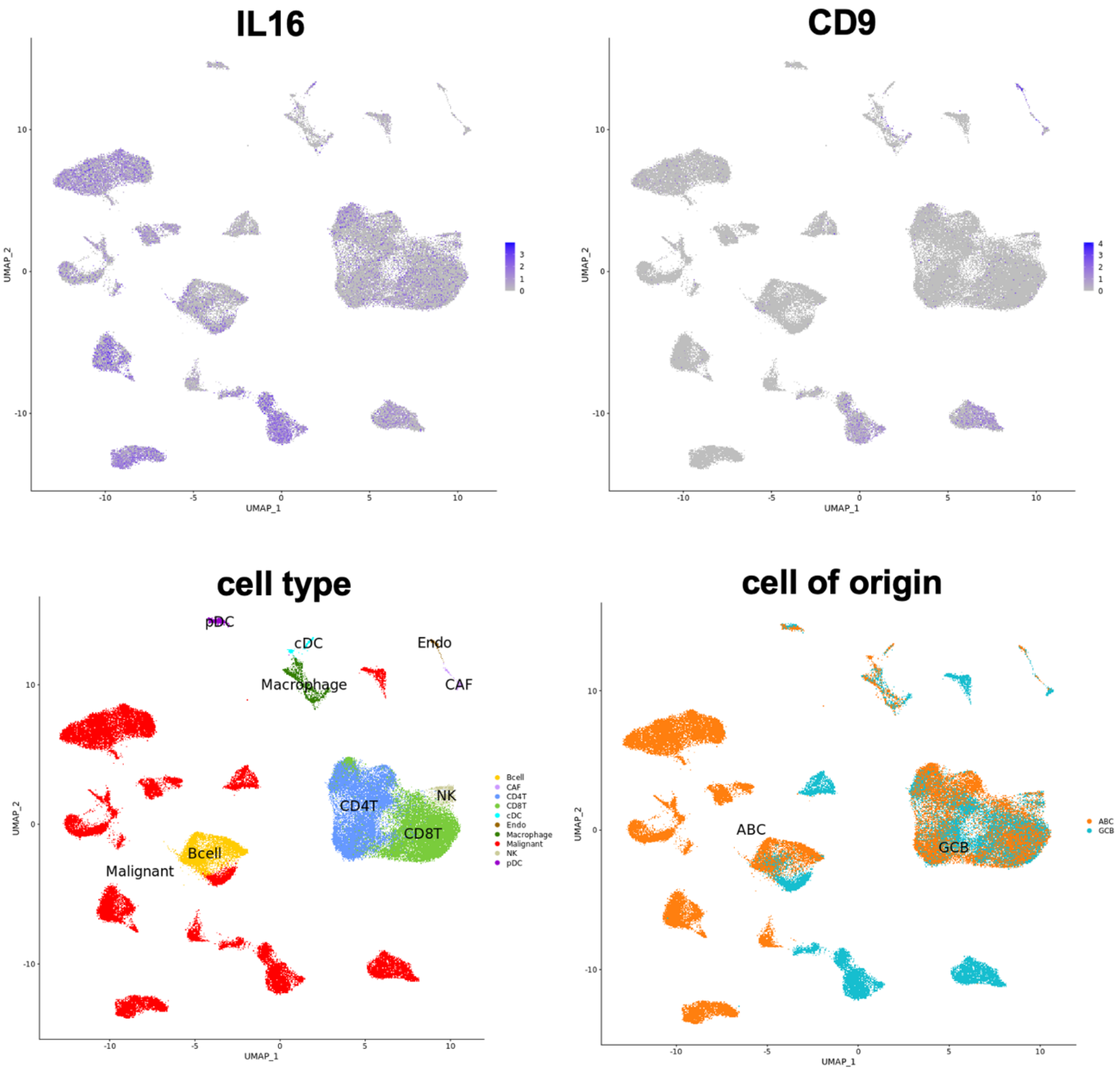
IL16
CD9
cell type
cell of origin
UMAP_1
UMAP_2
pDC
cDC
Endo
Macrophage
CAF
NK
CD4T
CD8T
Bcell
Malignant
Bcell
CAF
CD4T
CD8T
cDC
Endo
Macrophage
Malignant
NK
pDC
ABC
GCB
ABC
GCB

**Figure S19.** (A) Immunoblotting was performed to measure protein levels upon exposure to CD9-binding peptide (CD9-BP, 10µM), recombinant IL-16 (30ng/mL), the combination of both, or PBS (control); in VL51 parental and ibrutinib-resistant clones for NFkB1, NFkB2, and pAKT/AKT in two independent experiments. Vinculin was used as a loading control. (B) Protein quantification was done by normalizing first to vinculin and then to parental levels. The barplot represents the mean of the two independent experiments, parental in grey, resistant in orange. Error bars represent the standard deviation of the mean. Values at the end of the bars represent the p-value from a Welch Two Sample test.

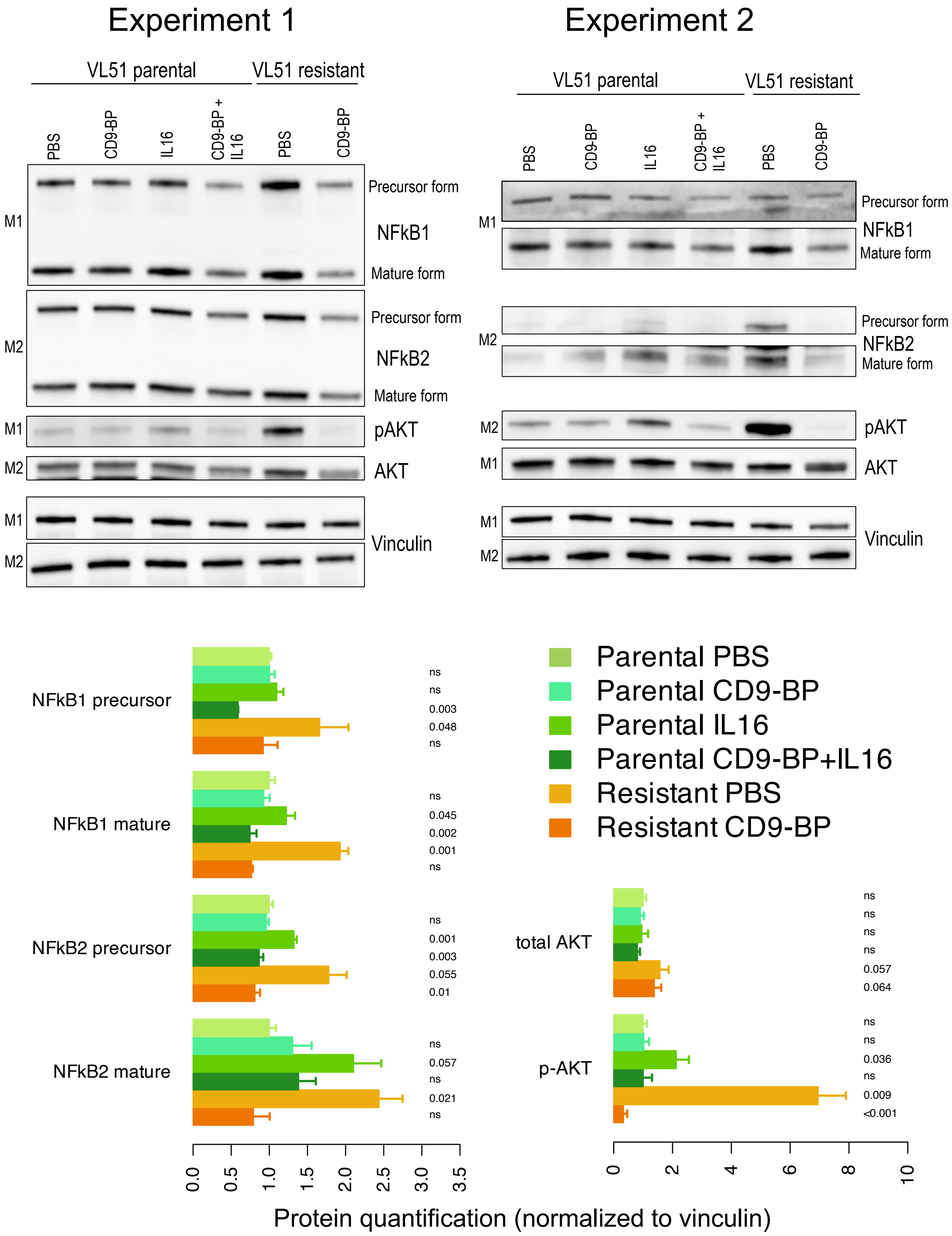

**Figure S20.** (A) Heatmap showing chromatin differential accessibility in consensus peaks based on ATAC-seq data in VL51 parental and ibrutinib-resistant cells. (B) VL51 ibrutinib-resistant Up and Down signatures were enriched in more accessible and less accessible peaks by gene-set enrichment analyses on ATAC-seq data. (C) FLI1 gene expression in VL51 ibrutinib-resistant (orange) and parental (grey) cells by RNA-sequencing. (D) Gene set enrichment analyses were performed using RNA-sequencing data on FLI1-regulated targets to compare ibrutinib-resistant (red) versus parental (blue) VL51 cells. (E) Ibrutinib sensitivity (IC50, log10) correlation (Pearson) with FLI1 expression (lcpm from RNA-seq) in ABC-DLBCL cell lines. (F) IL16-FLI1 correlation (Pearson, lcpm from RNA-seq) across B-cell lymphoma models (n=47). (G) Drug sensitivity to Chelerythrine, a FLI1 inhibitor [4], differed between parental (grey) and ibrutinib-resistant (orange) VL51 cells. (H) Synergy models of Chelerythrine and ibrutinib combo in VL51 parental (PAR, blue) and ibrutinib-resistant (IBR, red) cells. Drug sensitivity and synergy were evaluated by MTT assay (72 hours). Error bars correspond to the standard deviation of the mean; µM for micromolar. Data was derived from at least three independent experiments.

A

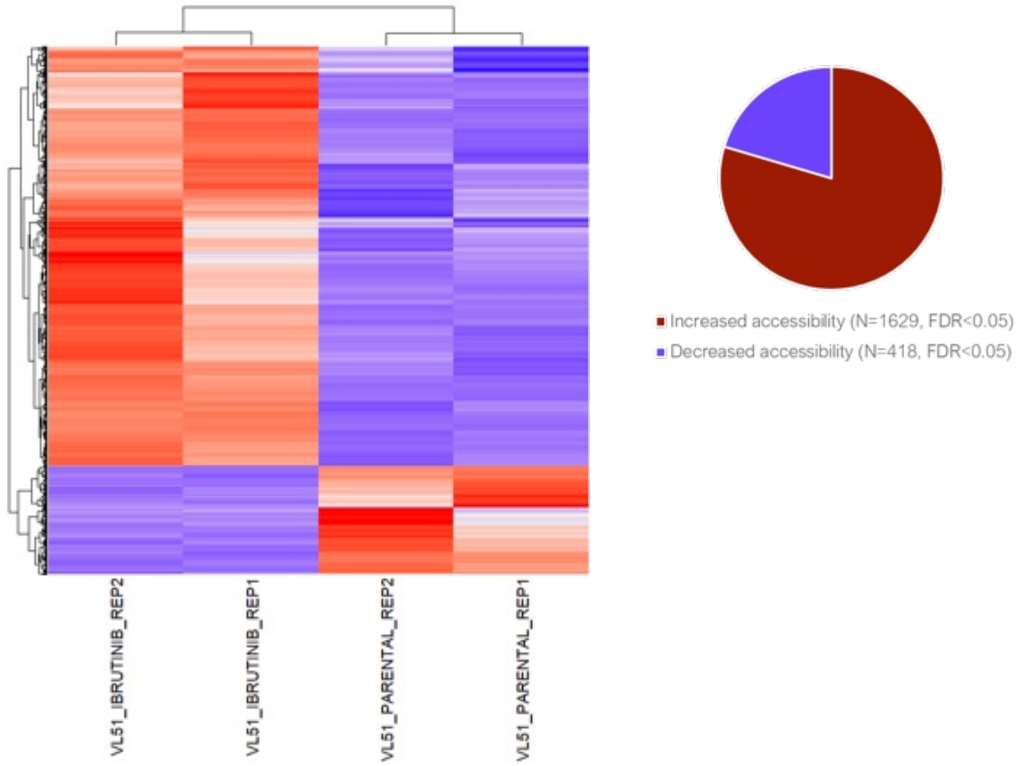


B

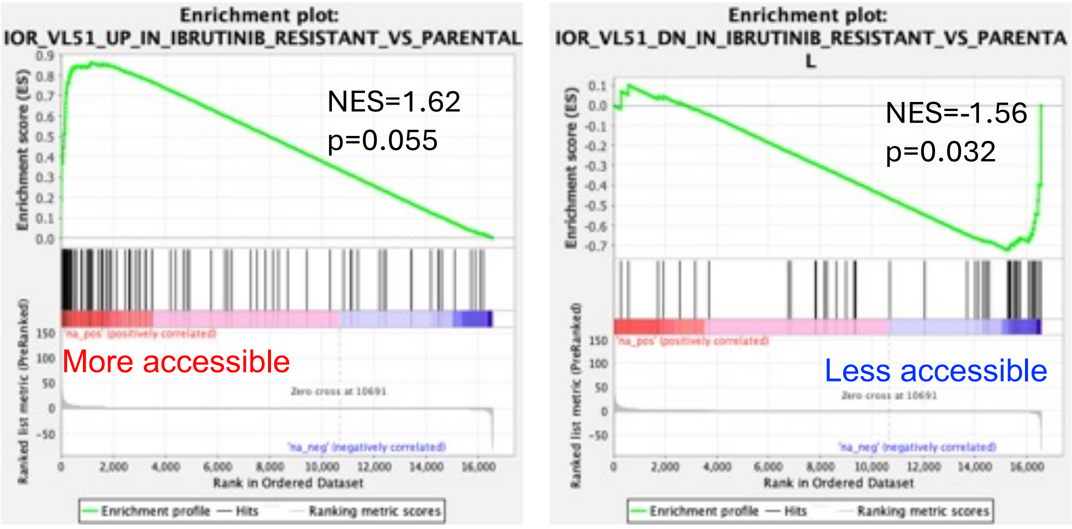

C

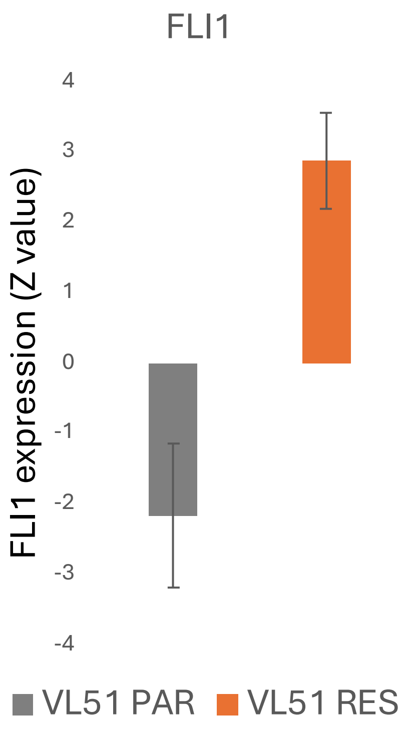

D

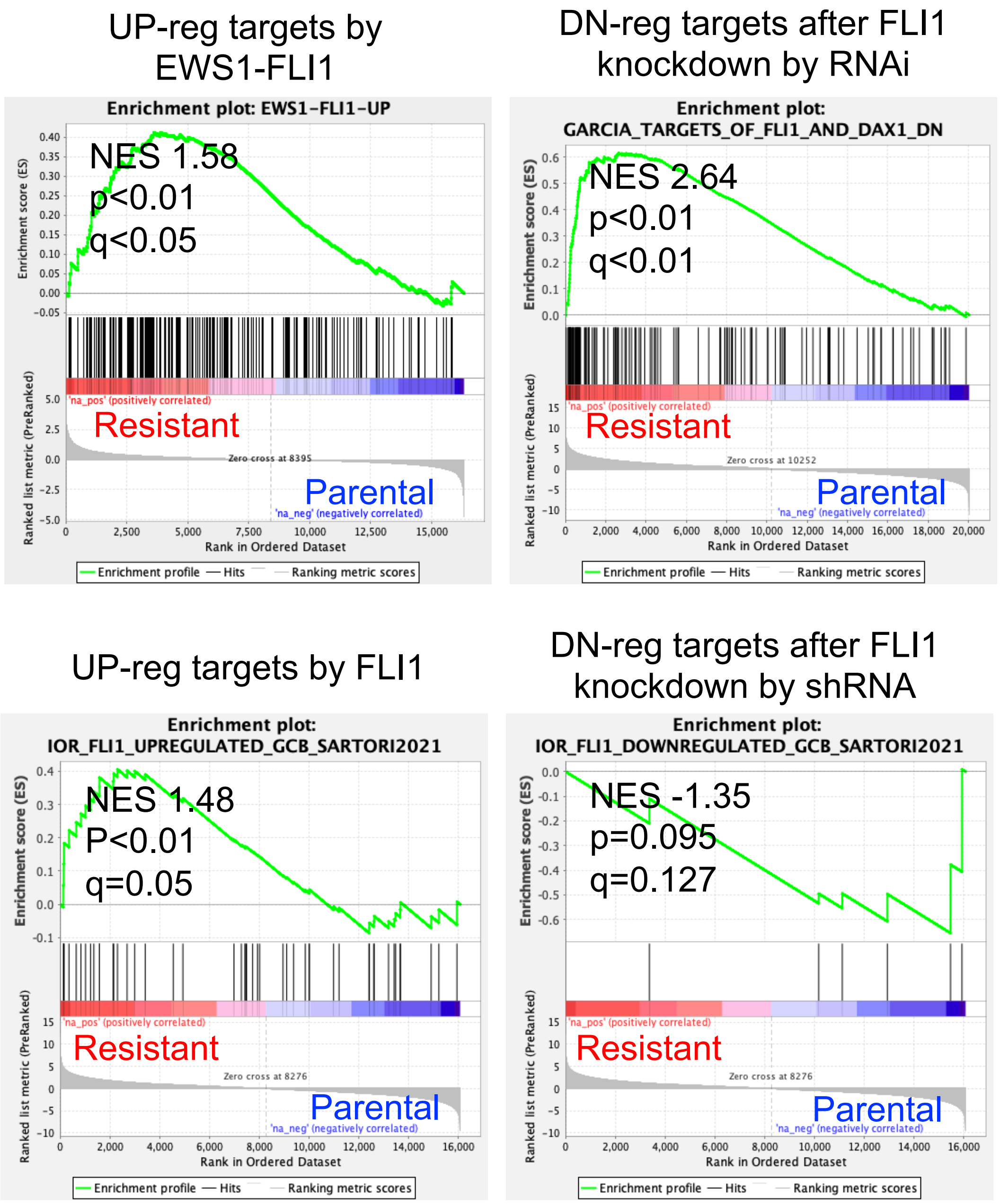

UP-reg targets by EWS1-FLI1
Enrichment plot: EWS1-FLI1-UP
NES 1.58
p<0.01
q<0.05
Resistant
Parental
'na_pos' (positively correlated)
'na_neg' (negatively correlated)
Zero cross at 8395
Enrichment score (ES)
Ranked list metric (PreRanked)
Rank in Ordered Dataset
Enrichment profile — Hits — Ranking metric scores
DN-reg targets after FLI1 knockdown by RNAi
Enrichment plot: GARCIA_TARGETS_OF_FLI1_AND_DAX1_DN
NES 2.64
p<0.01
q<0.01
Resistant
Parental
Zero cross at 10252
UP-reg targets by FLI1
Enrichment plot: IOR_FLI1_UPREGULATED_GCB_SARTORI2021
NES 1.48
P<0.01
q=0.05
Resistant
Parental
Zero cross at 8276
DN-reg targets after FLI1 knockdown by shRNA
Enrichment plot: IOR_FLI1_DOWNREGULATED_GCB_SARTORI2021
NES -1.35
p=0.095
q=0.127
Resistant
Parental
Zero cross at 8276

E

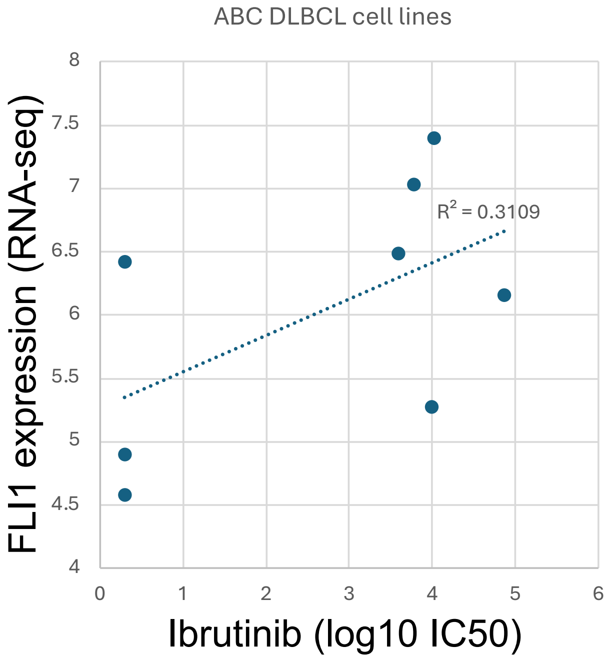
ABC DLBCL cell lines
R² = 0.3109
FLI1 expression (RNA-seq)
Ibrutinib (log10 IC50)
8
7.5
7
6.5
6
5.5
5
4.5
4
0
1
2
3
4
5
6

F

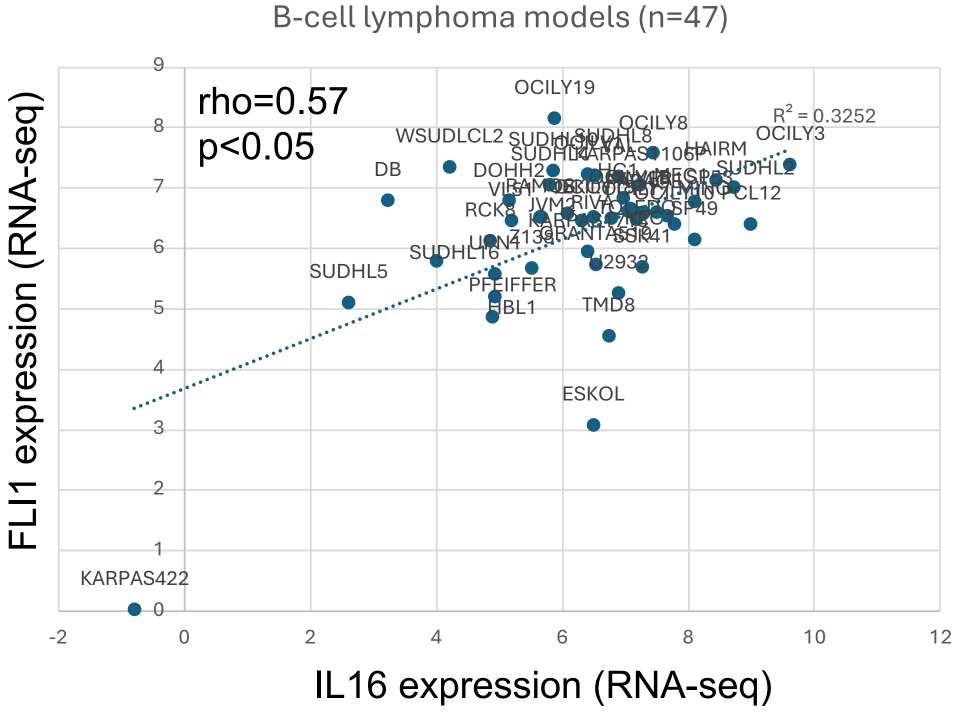
B-cell lymphoma models (n=47)
rho=0.57
p<0.05
R² = 0.3252
OCILY19
OCILY8
OCILY3
WSUDLCL2
HAIRM
SUDHL2
DB
DOHH2
PCL12
RCK8
SUDHL6
SUDHL5
PFEIFFER
HBL1
TMD8
ESKOL
KARPAS422
FLI1 expression (RNA-seq)
IL16 expression (RNA-seq)
9
8
7
6
5
4
3
2
1
0
-2
0
2
4
6
8
10
12

G

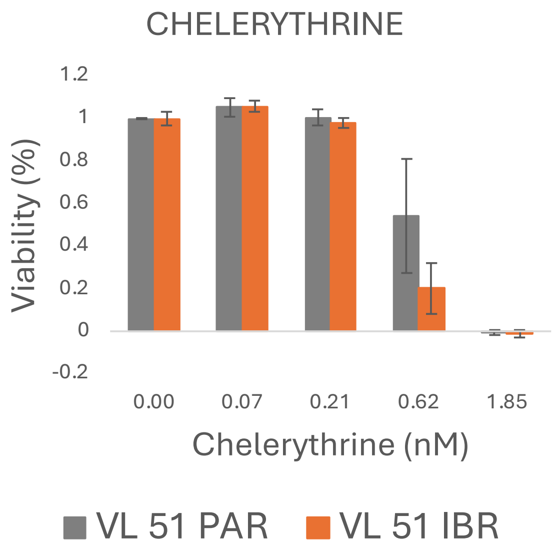
CHELERYTHRINE
Viability (%)
1.2
1
0.8
0.6
0.4
0.2
0
-0.2
0.00
0.07
0.21
0.62
1.85
Chelerythrine (nM)
VL 51 PAR
VL 51 IBR

H

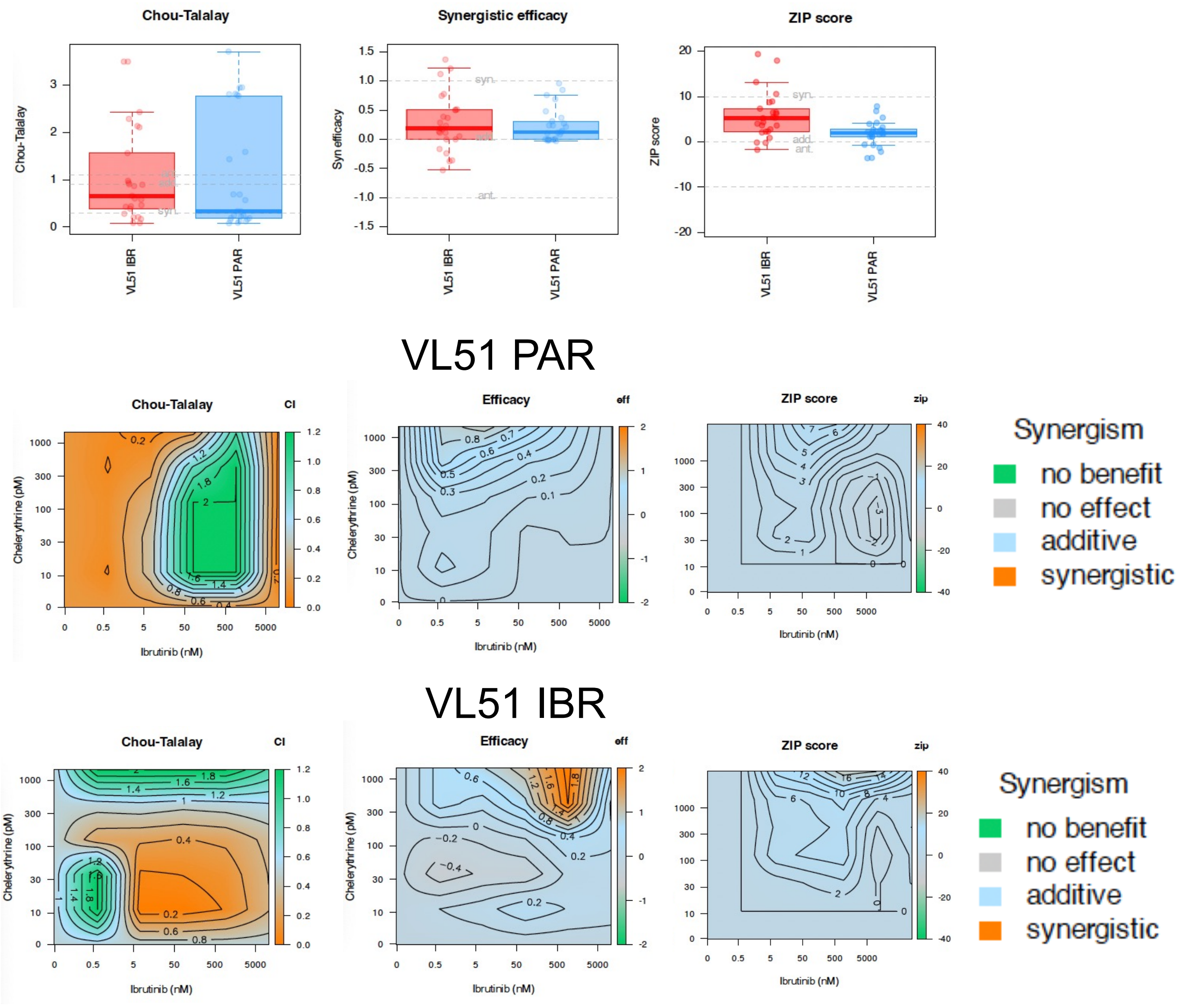
Chou-Talalay
Synergistic efficacy
ZIP score
Chou-Talalay
Syn efficacy
ZIP score
VL51 IBR
VL51 PAR
syn.
add.
ant.
VL51 PAR
Chou-Talalay
CI
Efficacy
eff
ZIP score
zip
Chelerythrine (pM)
Ibrutinib (nM)
Synergism
no benefit
no effect
additive
synergistic
VL51 IBR
Chou-Talalay
CI
Efficacy
eff
ZIP score
zip
Chelerythrine (pM)
Ibrutinib (nM)
Synergism
no benefit
no effect
additive
synergistic
